# Can the Heartbeat-Evoked Potential (HEP) be Separated from Cardiac Artefact (CA) Using Beamforming?

**DOI:** 10.64898/2026.07.11.737958

**Authors:** Rania-Iman Virjee, Rohan Kandasamy, Sarah N Garfinkel, Mahinda Yogarajah, Vladimir Litvak, David W Carmichael

**Author notes:** **Corresponding author:** Rania-Iman Virjee,. Joint senior authors (Vladimir Litvak, David W Carmichael).

## Abstract

The heartbeat-evoked potential (HEP), a cortical response to heartbeats and neural marker of interoception, is clinically relevant but heavily contaminated by scalp cardiac-artefact (CA), making reliable distinction challenging. Because HEP and cardiac potentials are anatomically distinct, they may be separable via beamforming, a source localisation method isolating brain activity while suppressing external interference. Here, the first known validation of EEG beamforming for HEP source reconstruction was attempted, quantifying waveform recovery and spatial accuracy using simulated data.

Linearly constrained minimal variance (LCMV) beamforming was used (A) to test signal recovery, 128-channel EEG was simulated for three models with known HEP waveforms: (1) a single right insula (R-Ins) HEP; (2) temporally distinct HEPs in R-Ins and right anterior cingulate cortex (R-ACC); (3) overlapping HEPs in both regions. Recovery was assessed by correlating reconstructed and known waveforms. (B) To test CA suppression, CA from brain-dead isoelectric EEGs was integrated to simulations, with varying source (-10-50dB) and sensor (0-30dB) signal-to-noise ratios (SNR). (C) To analyse empirical EEG from hypertensive and anxiety individuals (n=106). Null-space projection, using subject-specific QRS waveforms and its temporal derivative, removed residual CA in source waveforms.

In model-1, beamforming achieved near-perfect recovery without CA (r>0.99, error=0mm) at -30dB source SNR, remaining robust in the presence of CA (r=0.72-0.94, error=0-5.7mm) but degrading below -20dB (r<0.3, error=27mm). Models 2 and 3 showed similar recovery but introduced R-ACC-to-R-Ins leakage. In empirical data, beamforming revealed significant HEP activity in R-Ins and R-ACC (p<0.001). A significant late R-ACC HEP difference (250-500ms post-R-peak) between hypertension and controls emerged after excluding low-SNR subjects and strengthened following QRS cleaning (p=0.035, d=0.96).

LCMV beamforming reliably recovers simulated HEPs while suppressing CA given sufficient SNR. Applied empirically, beamforming detected clinically meaningful group differences in interoceptive regions (R-Ins/R-ACC), offering a source-level approach to separate HEP from CA that complements sensor-level methods.

## 1. Introduction

The heartbeat evoked potential (HEP), first described by Schandry, Sparrer and Weitkunat (1986), is a change in brain activity that is time and phase locked to each individual heartbeat. Generated with each cardiac cycle, the HEP has since been proposed as an implicit electrophysiological marker of interoception (Pollatos and Schandry, 2004). Interoception is defined as the process by which the nervous system senses, interprets, and integrates signals originating from within the body (e.g. hunger, thirst, cardiac and respiratory signals), providing a moment-by-moment mapping of the body’s internal landscape across conscious and unconscious levels (Khalsa et al., 2018). Heart-brain communication is bidirectional and cyclical, shaping both cognitive and emotional processes continuously (Park and Blanke, 2019). Since its initial characterisation, the HEP has been shown to be modulated by attention and arousal, and is detectable even without directed attention to the heartbeat (e.g. during sleep or at rest) (Pang et al., 2019; Petzsehner et al., 2019).

The HEP has emerged as a meaningful clinical measure, exhibiting both state and trait variation (Suksasilp and Garfinkel, 2022). Trait-level variation in HEP amplitude has been reported across psychiatric conditions associated with interoceptive dysfunction (i.e. anxiety disorders, borderline personality disorder) (Flasbeck et al., 2020; Pang et al., 2019). At the state level, HEP differences have been observed during interoceptive processing and in the periods preceding functional seizures (Desmedt et al., 2023; Elkommos et al., 2023; Hodossy et al., 2021; Kandasamy et al., 2026). Source localisation studies have consistently identified HEP generators in the insula and anterior cingulate cortex (Babo-Rebelo et al., 2016b, 2016a; Canales-Johnson et al., 2015; Park et al., 2014), somatosensory cortex and amygdala (Coll et al., 2021; H. Park and Blanke, 2019).

The HEP is typically measured with electroencephalography (EEG) or magnetoencephalography (MEG) time locked to the R-peaks or T waves of the individual’s electrocardiography (ECG). This produces a scalp-recorded event-related potential (ERP) typically visible between 200 to 600 ms post R peak (Park and Blanke, 2019). A fundamental and inherent challenge in HEP research is the cardiac artefact (CA), the electrical field generated by the electrophysiological activity of the heart spreading throughout the whole body, including the scalp termed the cardiac field artefact (CFA) (Arnau et al., 2023; Dirlich et al., 1997) as well as the pulsation-associated movement artefact (the cardioballistic artefact). The CA is itself time-locked to the R-peak. Thus, it cannot be separated from HEP by temporal averaging alone. The HEP is a low-amplitude transient neural response, typically in the range of a few microvolts (1-5 μV). In contrast, CA can be up to an order of magnitude larger (±10 μV). Critically, the CA reaches its maximum amplitude during the QRS complex and T-wave, with the latter directly overlapping with the temporal window of the HEP (Dirlich et al., 1997). This overlap inherently constrains the effectiveness of conventional ERP-style subtraction approaches. The CA is a dynamic artefact whose amplitude and morphology vary with factors including neck thickness, posture, heart rate, and respiration phase amongst others (Dirlich et al., 1997; Kern et al., 2013; Virjee et al., 2026). Because both CA and HEP are synchronised to the cardiac cycle, any R-peak locked average will contain a superposition of cortical HEP and CA. Standard sensor-level approaches to CA removal, in particular ICA-based methods, carry the risk of removing genuine HEP signal alongside artefact, due to the time-locked nature of the same event (Abolfathi and Mohebbi, 2024; Buot et al., 2021; Petzschner et al., 2019). This methodological challenge is reflected in the significant heterogeneity across HEP studies in terms of preprocessing choices, and time windows (Coll et al., 2021; Virjee et al., 2026). With the growing interest in HEP, as a clinical and scientific measure, there is a pressing need for robust, principled, and standardised methodologies for its measurement. In this context, source reconstruction that moves HEP analysis into brain space, where CA should be suppressed due to its distinct spatial origins, offers a principled solution to this problem.

Linearly Constrained Minimum Variance (LCMV) beamforming is a spatial filtering technique originally developed for sonar and radar applications in the early years of the 20^th^ century (Van Veen et al., 1997; Van Veen and Buckley, 1988) before being utilised in neuroscience as a versatile and robust tool and as a technique to interpret the neural basis of EEG and MEG data (Westner *et al*., 2022). LCMV beamforming, as proposed by Van Veen et al. (1997), operates by computing, for every source location *r*, a set of spatial weights, such that the activity originating from *r* is passed with unit gain to preserve the source of interest while minimising output variance to suppress contributions from other locations and noise. This is achieved without requiring *a priori* assumptions about the number of active sources, making LCMV beamforming a data-driven method well suited for exploratory and clinical applications. Its spatial filtering properties make it well-suited to the HEP problem: by computing location-specific weights that pass activity from a target source while suppressing contributions from other locations, the beamformer can in principle isolate cortical HEP activity from the spatially distinct CA.

To the best of our knowledge, LCMV beamforming has not previously been applied to HEP extraction. Validating this approach requires addressing several fundamental questions: how accurately does beamforming recover HEP source waveforms under realistic noise conditions? Does the presence of CA substantially impair beamformer performance, or can the spatial filter tolerate and suppress cardiac artefact contamination? These questions are addressed here through a two-part study. First, a simulation study evaluated LCMV beamforming performance across a fully factorial design spanning three HEP source models of increasing complexity, combined with source-space noise, sensor-space noise, and empirically derived CA obtained from isoelectric EEG of brain-dead individuals. Second, the validated LCMV beamforming pipeline was applied to retrospective empirical data to investigate whether LCMV beamforming yields anatomically plausible HEP waveforms and is sensitive to clinically meaningful group differences.

This study offers the first proof-of-concept of EEG beamforming for HEP source reconstruction, aiming to quantify source waveform recovery and spatial localisation accuracy.

## 2. Methods

### 2.1. Overview of proof of concept

The methods are organised in two parts. First, a proof-of-concept simulation study was conducted to validate the LCMV beamforming pipeline under controlled conditions with a known source. Second, the validated pipeline was then applied to empirical clinical EEG data. Both analyses employ the same beamforming framework as detailed below.

### 2.2. LCMV beamforming

Beamforming is a source reconstruction technique acting as a spatial filter that reconstructs neural source activity from sensor level EEG or MEG data. It therefore acts as a bridge between two spatial domains (Westner *et al*., 2022): source space and sensor space. Source space refers to the true locations and orientations of focal brain sources. Sensor space constitutes signals that are directly recorded from the scalp such as with EEG or MEG. In essence, beamforming operates by sweeping through a defined set of source locations, calculating weights for each source location, the spatial filter, and applying the weights to the sensor space measurements. The weights are computed to be able to reconstruct source activity while suppressing contributions from other brain regions and noise. The output will be the beamformer source space activity for each location (Westner *et al*., 2022). Therefore, the spatial filter represents the contribution of each scalp sensor to each source estimate location (Baillet et al., 2001; Van Veen et al., 1997; Westner et al., 2022).

As explained in Brookes *et al*. (2008), Beamforming estimates the source activity at a source location *r* and time *t*, using a weighted sum of the electrical potential measurements recorded at each of the *n* EEG electrodes (sensor signals):

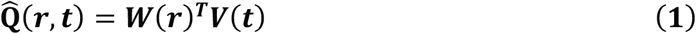

Where:

- Q̂(***r***, ***t***) is the 3-dimensional source amplitude vector 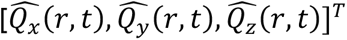 at location *r* and time *t*
- ***W***(***r***) is a 3 x *n* weight matrix specifically for source location *r* with one weight vector per orientation *x*, *y* and *z*. The spatial filter ***W***(***r***) will be unique for each source location. The beamformer weights are therefore location-specific to extract the source at that location while suppressing activity from other locations.
- ***V***(***t***) are the EEG electric potential measurements expressed as an *n* dimensional column vector at time *t*

Two components are essential to compute the spatial filter: the forward model and the data covariance matrix.

The forward model or lead field, *L*(*r*), of dimension 3 *x n*, describes the mapping from a source location *r* to the recorded EEG scalp measurements. The three columns of the lead field encode the electrical potential measured at each of the *n* electrodes in response to a unit-amplitude source at location *r*. Each column corresponds to a dipole orientated in the *x*, *y and z* directions respectively (Brookes *et al*., 2008). It is computed from the geometry and conductivity of the head using a boundary element method (BEM). The covariance matrix, *C*, describes how the EEG electrodes co-vary in time and is denoted as:

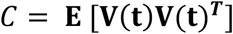

Thus, the covariance matrix captures the electrodes that are “seen” by the same brain sources. High covariance values between electrodes will indicate that they are sensitive to the same underlying sources. This structure is utilised by the beamformer to suppress shared interference while preserving the target source.

#### The LCMV solution

Linearly Constrained Minimum Variance (LCMV) beamformer is founded on the principle of computing beamformer weights that minimise the total output variance of the signal

Q̂(***r***, ***t***) while ensuring that activity occurring at source location *r*, remains in Q̂***r***, ***t***). This translates to (Brookes *et al*., 2008):

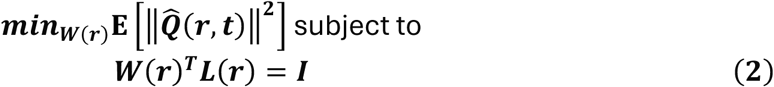

Where:

- ***I*** is the identity matrix
- the expression ***W***(***r***)***^T^L***(***r***) allows to meet the unity unit gain constraint for the spatial filter, i.e. activity at source location *r* is preserved
- 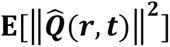 ensures minimum variance, i.e. suppressing all other sources and noise other than the source at location *r*

The closed-form solution is:

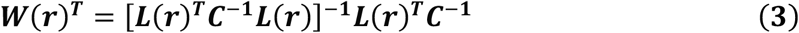

Therefore, the beamformer weights depend on the inverse of the data covariance matrix and the forward solution at location *r*. The beamformer uses ***C***^−**1**^ to identify and down-weight sensor combinations that carry variance from other brain locations, while ***L***(***r***) ensures the filter is tuned to the specific spatial signature of the target source.

Importantly, LCMV beamforming assumes that brain sources are temporally uncorrelated. If two sources are correlated, beamforming will suppress one or both, as their shared variance will be interpreted as a single source. This assumption shaped the simulation design described in section 2.3 where two HEP sources were modelled simultaneously and orthogonalized to prevent correlation-driven suppression.

#### Source power and regularisation

The beamformer projected power can then be defined as:

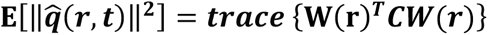

The trace operation sums the estimated power across all three dipole orientations. This provides the power of the reconstructed source signal at location r and is invariant to the orientation of the underlying neural source.

In practice, ***C*** is estimated from finite data and may be rank-deficient or ill-conditioned. This can occur in high-density EEG where the number of electrodes approaches or exceeds the number of independent data samples. An ill-conditioned ***C*** produces an instable inverse, amplifying noise in the weights. Therefore, throughout this study, the covariance was regularised using principal component analysis (PCA) with dimensionality reduction to 100 components.

For a full review on the theoretical underpinnings of LCMV beamforming, please refer to Brookes et al., (2008) and Westner et al., (2022).

#### Common spatial filter strategy

When comparing source power between two conditions (here, a baseline window including the R peak and a HEP activation window) a common spatial filter strategy was used to prevent spurious differences arising from computing separate covariance matrices per condition because the spatial filter itself would differ between conditions. Consequently, a single covariance matrix was estimated from data equally pooled across both windows, and the resulting weights were then applied separately to each window to compute condition-specific power. Thus, differences in reconstructed source power can be attributed to genuine differences in activity rather that differences in the spatial filter.

### 2.3. Simulation study

In order to evaluate whether LCMV beamforming can reliably recover HEP in shape and time, in the presence of CA and without CA, and how its performance varies with SNR, synthetic EEG datasets were generated with known sources under a fully controlled factorial design.

#### 2.3.1. The forward model

Forward modelling was performed using the FreeSurfer ‘fsaverage’ template brain. A three-layer boundary element model (BEM) was constructed with standard EEG conductivities, 0.3 S/m, 0.0006 S/m, 0.3 S/m, representing the inner skull, outer skull, and skin respectively. The cortical source space was defined using an octahedral (oct6) subdivision, and sources within 5 mm of the inner skull were excluded. The forward solution was computed for a 128-channel EEG BioSemi electrode layout using MNE-Python (Python version 3.13.7, MNE version 1.10.1). Electrode positions and event markers were derived from an empirical BioSemi BDF (sample dataset available on the SPM website comprising of an auditory oddball paradigm). Auditory stimulus-onset markers were used as timing anchors, providing a realistic non-uniform event structure onto which the simulated HEP epochs were built, rather than genuine R-peak timings.

#### 2.3.2. HEP source models

Simulated HEP sources were placed in the right insula (R-Ins) and right rostral anterior cingulate cortex (R-ACC), based on prior interoceptive network studies (Babo-Rebelo et al., 2016a; Canales-Johnson et al., 2015; Coll et al., 2021; Park et al., 2014). A source amplitude of 2 *nAm* was chosen to match previously reported empirical and simulated HEP magnitudes (Coll et al., 2021). Regions of interest (ROIs) were defined using the Desikan-Killiany (aparc, Freesurfer) parcellation, selecting a 10 mm patch centred within each label. All dipoles within a ROI shared a common time course and were oriented normal to the cortical surface. All simulated datasets were generated using MNE python (Python version 3.13.7, MNE version 1.10.1, NumPy version 2.3.2) and MATLAB (version 25.1.0.2943329 R2025a). LCMV beamforming was implemented using the DAiSS (Data Analysis in Source Space) toolbox in SPM12 (Henson et al., 2019; Litvak et al., 2011).

Three source models of increasing complexity were constructed and differed in terms of number of ROIs, duration, timing, and relative amplitude (figure 1):

**Figure 1.**
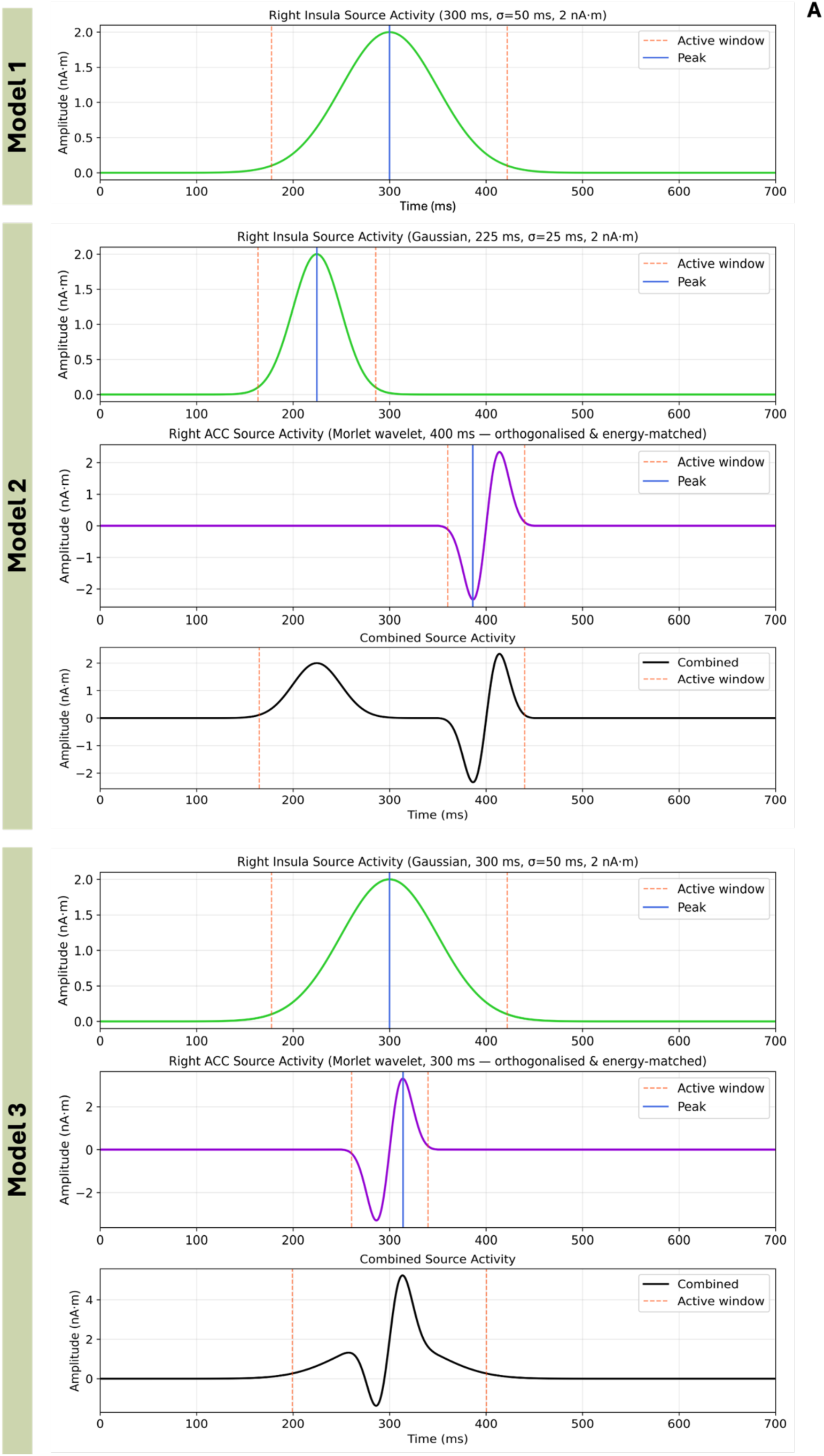
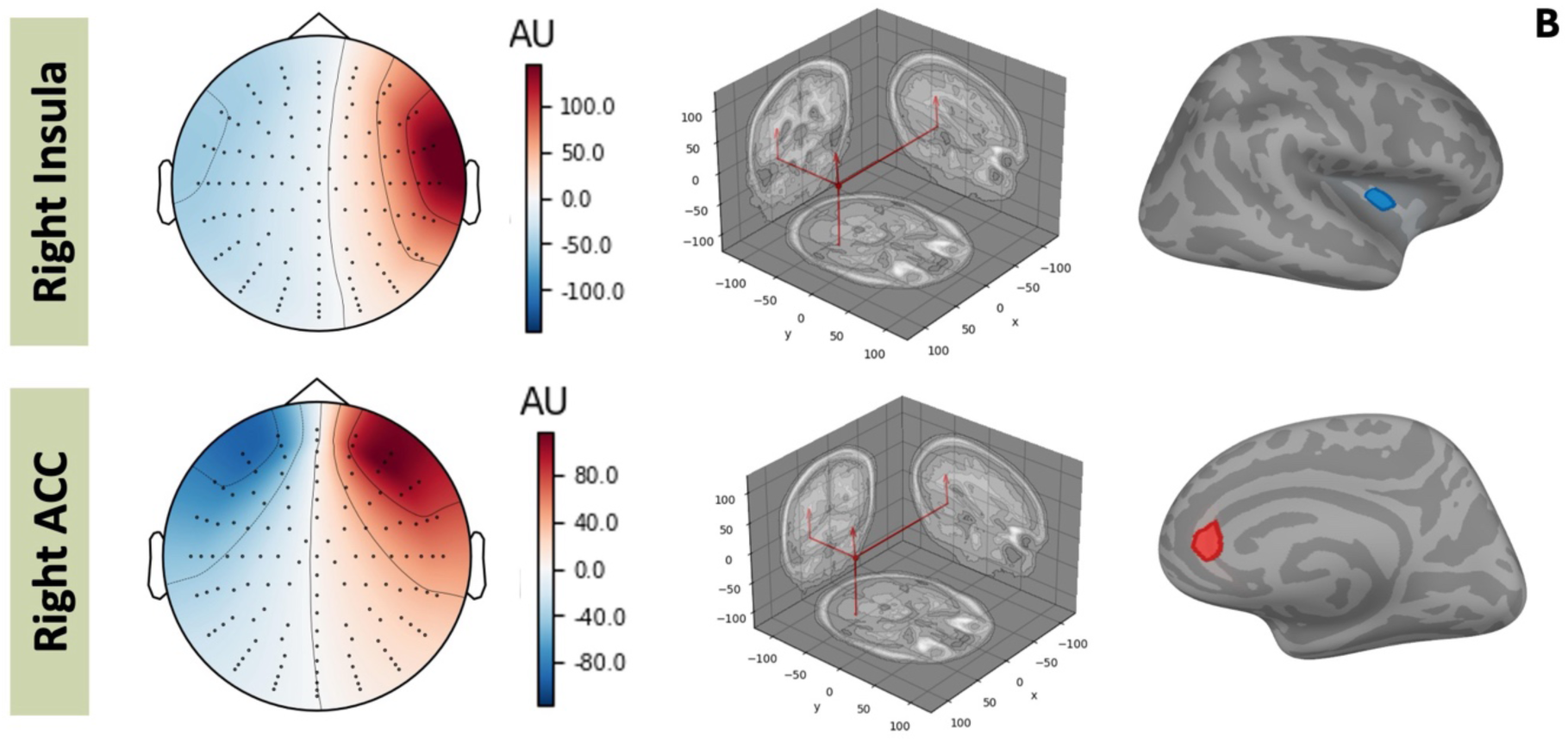
Schematic diagram representing the temporal profile of each model (A), and source localisation and sensor-level projections of the simulated sources in the R-Ins and R-ACC (B).

Model 1 contained a single source in R-Ins, modelled as a Gaussian waveform representing HEP (peak latency *μ* = 300 *ms*, *σ* = 50 *ms*, amplitude *A* = 2 *nAm*, 200 to 400 ms post R-peak), illustrating the simplest scenario for beamforming. A Gaussian waveform was chosen as it most closely aligns with waveform morphology reported in intracranial EEG recordings (Wang et al., 2025). Model 2 contained two sources active in non-overlapping windows: R-Ins active 150-300 ms post R-peak (*μ* = 225 *ms*, *σ* = 25 *ms*, amplitude *A* = 2 *nAm*), and R-ACC active 350-450 ms post R-peak (*μ* = 400 *ms*). This tests whether beamforming can resolve spatially distinct and temporally separated sources. Finally, model 3 contained the same two active sources as model 2, but sources were overlapping in time (200-400 ms post R-peak, *μ* = 300 *ms*). This is the most challenging model for source separation.

In model 3, using two Gaussian waveforms would yield highly correlated source time courses violating the beamforming assumption of temporal independence and making it difficult to estimate the leakage between sources. To circumvent this, the R-ACC source was modelled as the imaginary part of a single-cycle Morlet wavelet (centre-frequency 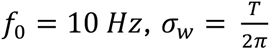, where *T* = 1⁄*f*_0_ and is the wavelet duration) and orthogonalized with respect to the R-Ins Gaussian waveform using Gram-Schmidt orthogonalization. Additionally, both waveforms were energy matched to assure equal signal power across both sources. This temporal independence ensured the evaluation of the beamformer was based only on the spatial properties of the sources.

Both sources were assigned a common dipole orientation defined as the mean surface normal across R-Ins dipoles. This parallel orientation was chosen as it produces maximally correlated scalp topographies and represents the most challenging spatial arrangement for the beamformer source separation. Furthermore, the lead field vectors for both locations will be nearly collinear. This means that the spatial filter, although intended to pass only one source, will inevitably pass both, resulting in source leakage. Source activity at one location bleeds into the reconstruction of the other.

#### 2.3.3. Noise and CA addition

To systematically evaluate beamformer robustness, noise and artefact were added in sequential stages (figure 3).

**Figure 2.**
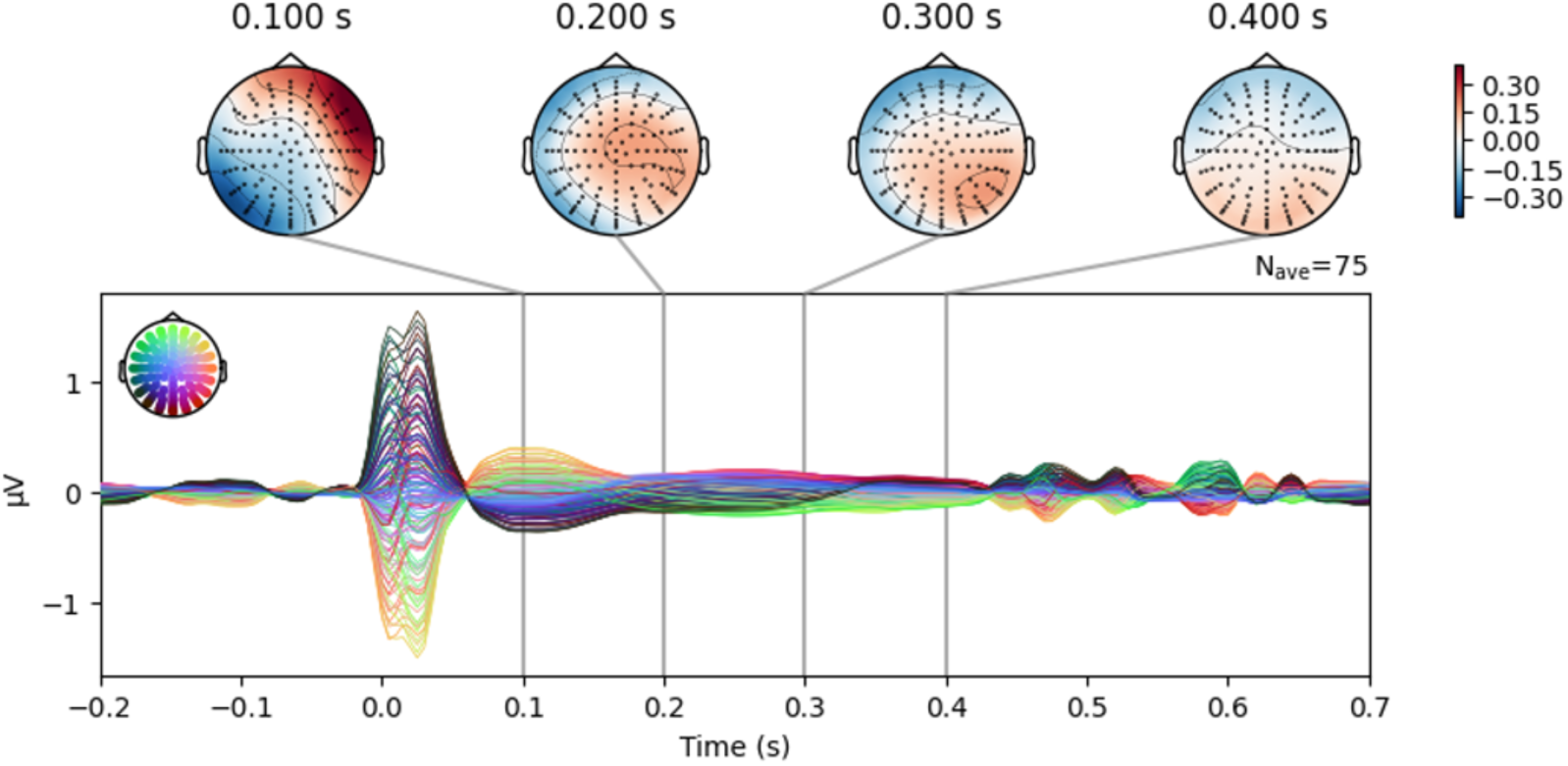
Grand-average empirical CA waveforms across retained epochs from brain-dead individuals (TUEG dataset), shown across all 128 channels. The CA appears to strongly coincide with the HEP window post R-peak.

**Figure 3.**
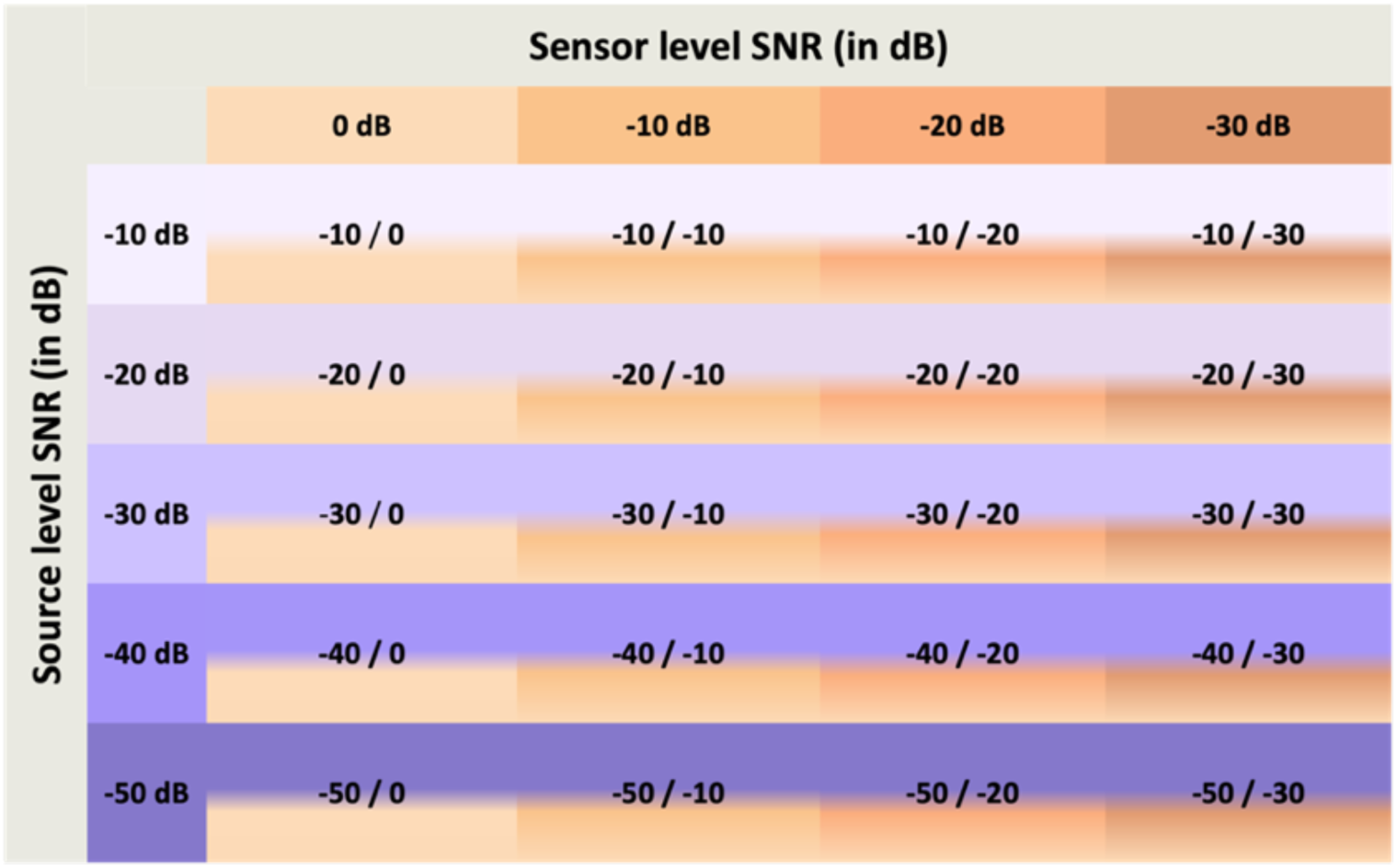
Matrix of source-space and sensor-space noise used for simulating EEG data.

First, source space noise was modelled. Spatiotemporally uncorrelated pink noise was added in source space prior to forward projection to model intrinsic background neural activity and non-cardiac physiological processes, at five SNR levels: -10 dB, -20 dB, -30 dB, -40 dB, -50 dB. Noise amplitude was calibrated per file relative to the root-mean-square (RMS) of the clean source signal following *RMS_noise_* = *RMS_signal_*⁄10*^SNRdB^*^⁄20^. Seventy-five unique independent noise realisations were generated per level per model. Second, Forward projection was computed. Each source-space dataset was projected through the forward model to sensor space, generating 128-channel EEG. Data were epoched -200 to +700 ms around R-peaks, baseline corrected -200 to 0 ms and resampled at 200 Hz. Third, sensor space noise was modelled. White Gaussian noise was added at four SNR levels (0 dB, -10 dB, -20 dB, -30 dB), and calibrated relative to the RMS of each individual file to preserve relative SNR across conditions. Fixed pseudorandom seeds ensured reproducibility. Lastly, Cardiac artefact (CA) was integrated. To derive an empirical CA, individuals who retain cardiac activity but have very limited brain activity may provide a model of CA in scalp EEG with very little cortical activity within the regions typically associated with HEP generation. To this end, EEG from 75 records classified as brain-dead were analysed from the Temple University EEG dataset – TUEG (Obeid & Picone, 2016). The EEG from each record was reviewed and confirmed as lacking visually detectable brain activity by an expert reviewer (RK). CA recordings were pre-processed as follows: bandpass 0.5-40 Hz, downsampling to 200 Hz, ICA to remove non-cardiac artefacts using MNE ICLabel (version 0.7.0), epoched -200 to + 700 ms, baseline corrected -200 to -50 ms, amplitude threshold 150 *μV*. As the recordings used a standard 10-20 montage (19 channels), missing channels were reconstructed using spherical-spline interpolation and reordered to the canonical BioSemi128 layout. Six-hundred CA epochs were retained per participant. Each of the previously generated 75 simulations instances was paired with a unique CA recording.

This pipeline generated for each model a 5 (source SNR levels) x 4 (sensor SNR levels) x 2 (with and without CA) design (figure 4, figure 5).

**Figure 4.**
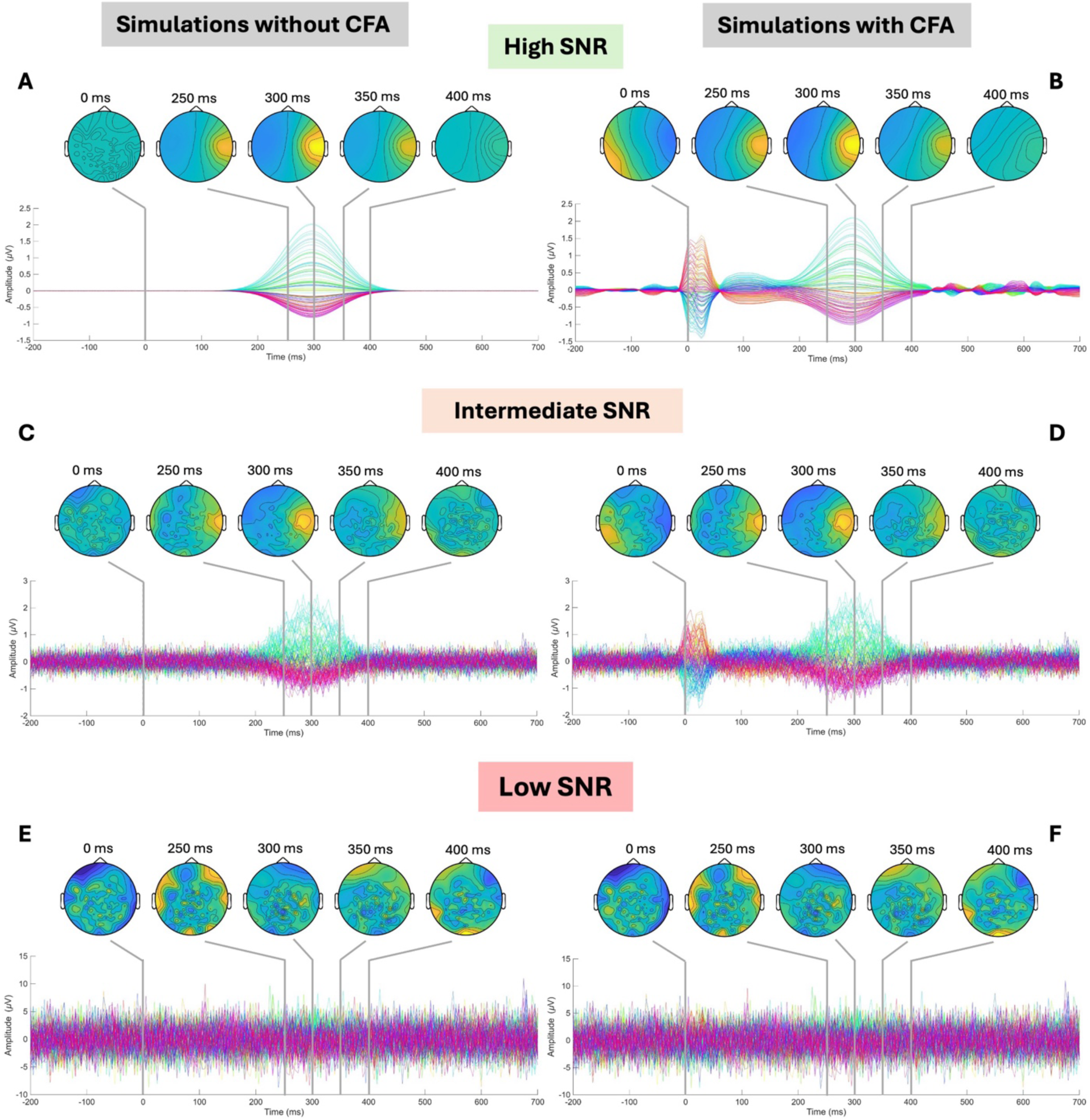
Model 1- simulation results without CA and in the presence of CA. Model 1 contained a single source in R-Ins, modelled as a Gaussian waveform representing HEP. Simulations without CA (A, C, E) and in the presence of CA (B, D, F) are presented here at varying SNR levels. High SNR corresponds to source SNR at -10 dB and sensor SNR at 0dB (A, B). Intermediate SNR corresponds to source SNR at -40 dB and sensor SNR at -20dB (C, D). Low SNR corresponds to source SNR at -50 dB and sensor SNR at -30dB (E, F). As SNR decreases, the HEP sources are drowned in noise at the scalp.

**Figure 5.**
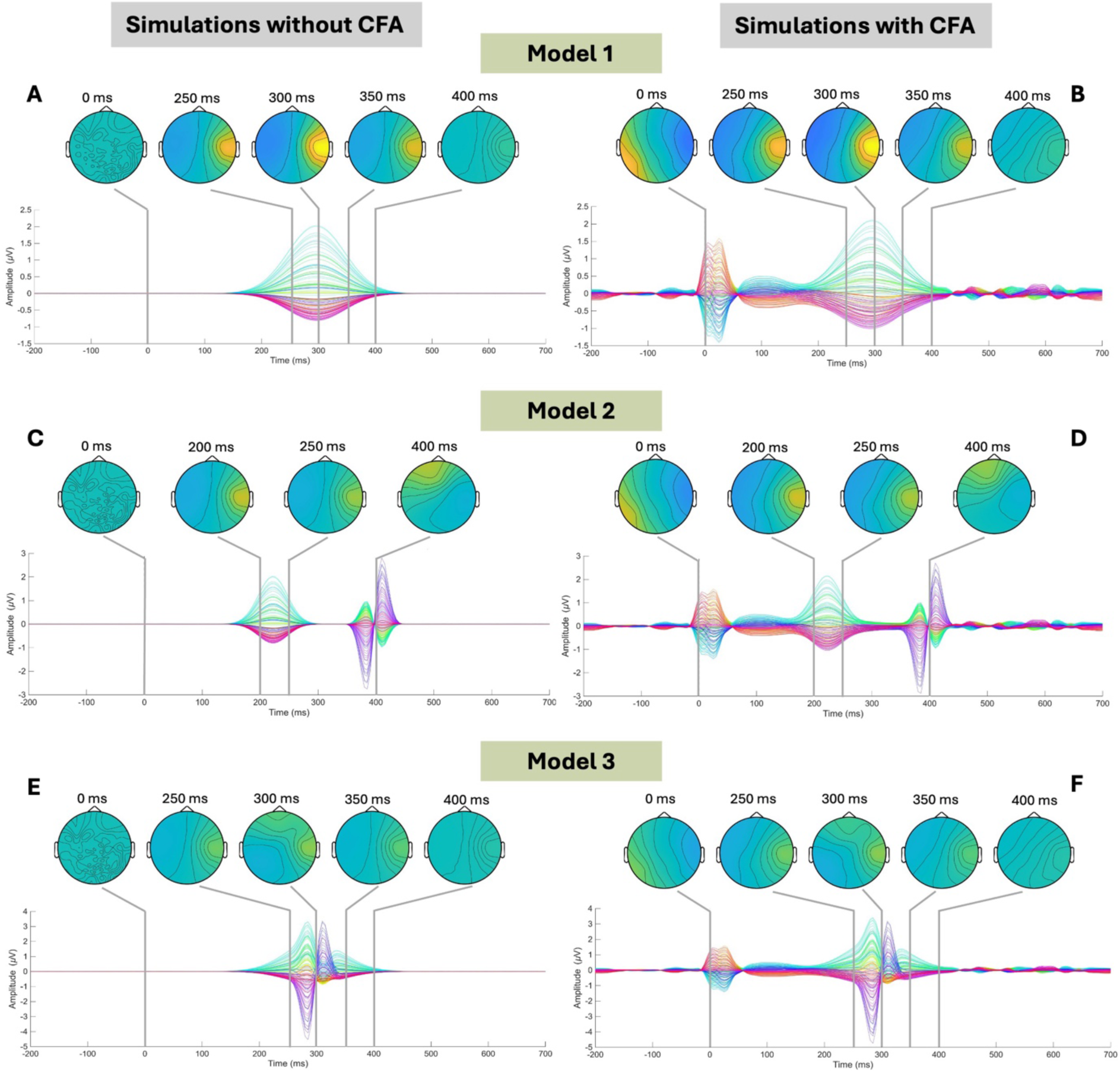
Simulation results without CA and in the presence of CA for each model at high SNR (source SNR at -10 dB and sensor SNR at 0dB). Model 1 (A, B) with a single source in R-Ins, modelled as a Gaussian waveform representing HEP, model 2 (C, D) with two temporally distinct HEPs (R-Ins and R-ACC), and model 3 (E, F) with two temporally overlapping HEPs in the same regions (R-Ins and R-ACC).

#### 2.3.4. Beamforming pipeline

The DAiSS beamforming toolbox, implemented within SPM12, was used to perform source reconstruction. The complete LCMV beamforming pipeline used here for source power imaging and source waveform reconstruction has been made available as example pipelines within DAiSS to support reproducibility and reuse for HEP source analysis.

For source power imaging, a volumetric source space was defined as a 3D grid with 5mm resolution in Montreal Neurological Institute (MNI) space, constrained to the inner skull. The sensor covariance matrix was estimated over -100 to + 400 ms (0-30 Hz) using the common spatial filter approach. PCA regularisation (100 dimensions) was applied. LCMV weights were computed per voxel and applied to estimate source power separately in baseline (-100 to +100 ms) and activation (200 to 400 ms) windows. Critically, unlike in most beamformer applications, the weights were applied to the average cardiac evoked response rather than single trials. This is because the HEP is a cardiac-locked average response, where single heartbeats are dominated by non-time-locked ongoing activity that is not part of the signal of interest. A differential contrast image (activation minus baseline) was computed and written to MNI space and adapted to the baseline and activation windows for each model.

To reconstruct time-resolved source waveforms, a VOI-based approach was applied to R-Ins (MNI: 36, 0, 0) and R-ACC (MNI: 6, 40, 10), each defined as a 10 mm radius sphere sampled at 5 mm resolution. MNI coordinates were identified from a noise-free simulation. The covariance was estimated across the full epoch (0-30 Hz, PCA to 100 dimensions). The voxel with maximum power within each VOI was selected, its spatial filter was applied to the average cardiac evoked response to produce an evoked source waveform per VOI.

#### 2.3.5. Validation metrics

Beamformer performance was quantified using two metrics computed against the known input source for each simulated file: waveform recovery and localisation error. Waveform recovery was expressed using the absolute Pearson correlation |*r*| between the reconstructed source time course and the known Gaussian waveform over the full time series, with absolute value taken to account for the sign ambiguity inherent to LCMV beamforming. For localisation error, this was the Euclidian distance (mm) between the peak voxel of the contrast image and the true simulated dipole location.

### 2.4. Applying beamforming to empirical clinical EEG data

#### 2.4.1. Participants and sensor-space preprocessing

The 128-channel EEG datasets from (Yoris et al., 2018, 2017) comprised of individuals with hypertension (HTN), healthy controls (NHTN), individuals with anxiety including panic and OCD (ANX), and healthy controls (NANX). Data was provided pre-processed: down-sampled to 256 Hz and band-passed filtered (0.3-50 Hz). Raw data were not available.

ECG quality was assessed for each participant by visual inspection of R-peak detection plots (NeuroKit2 library, Makowski et al., 2021). Individuals with invalid ECG channels were excluded, leaving 22 HTN, 24 NHTN, 22 ANX, and 38 NANX subjects remaining for further analysis.

All channel names were standardised to the BioSemi 128-channel naming convention, and the corresponding montage was applied. Noisy channels were identified and interpolated with PyPREP (version 0.5.0). Data were averaged-referenced and decomposed using extended Infomax ICA (48 components). ICA components were automatically labelled using ICLabel on 1-100 Hz filtered data, then reviewed manually. Non-neural components were removed, retaining heartbeat-associated components to avoid inadvertently removing HEP from CA. R-peaks were detected using NeuroKit2 with an adaptive outlier rejection and missing peaks were interpolated at the local median RR interval (0.7 s). Epochs were baseline corrected (-200 ms to -40 ms pre-R-peak).

#### 2.4.2. Beamforming pipeline

Prior to beamforming, bad channels were identified and excluded rather than being interpolated. This is because interpolation does not add new information but introduces linear dependencies that can compromise the beamformer solution. Data were bandpass filtered (0.5-30 Hz), average-referenced, and epoched using the same R-peak detection, outlier rejection, and baseline correction (-200 ms to -40 ms) as previously described (section 2.4.1). Power images were not computed as SNR was too low to yield meaningful results (based on our simulations).

#### 2.4.3. Beamforming and QRS cleaning

The same DAiSS/SPM12 beamforming pipeline described in section 2.3.4. was applied. However, the VOI sphere for source extraction was extended to 20 mm radius to accommodate anatomical variability across participants.

Null-space projection was applied to remove residual CA in the reconstructed source waveforms (figure 6). This was also applied to scalp data. For this, a subject-specific ECG template was constructed by creating epochs around R-peaks (-100 to 500 ms) and averaging across all beats. This window was chosen to capture the full QRS complex and T-wave morphology to ensure the HEP detected in source waveforms was not mixed with residual CA. The template and its temporal derivative were normalised to unit norm and concatenated as a two-column basis matrix spanning the CA subspace and making it possible to compensate for small temporal shifts between the ECG and the virtual electrodes. A null projector *P* = *I* − *UU^T^*, where U contains the left singular vectors of the basis matrix, was then applied to the average source waveforms, removing any component lying within the CA subspace while leaving orthogonal signal intact.

**Figure 6.**
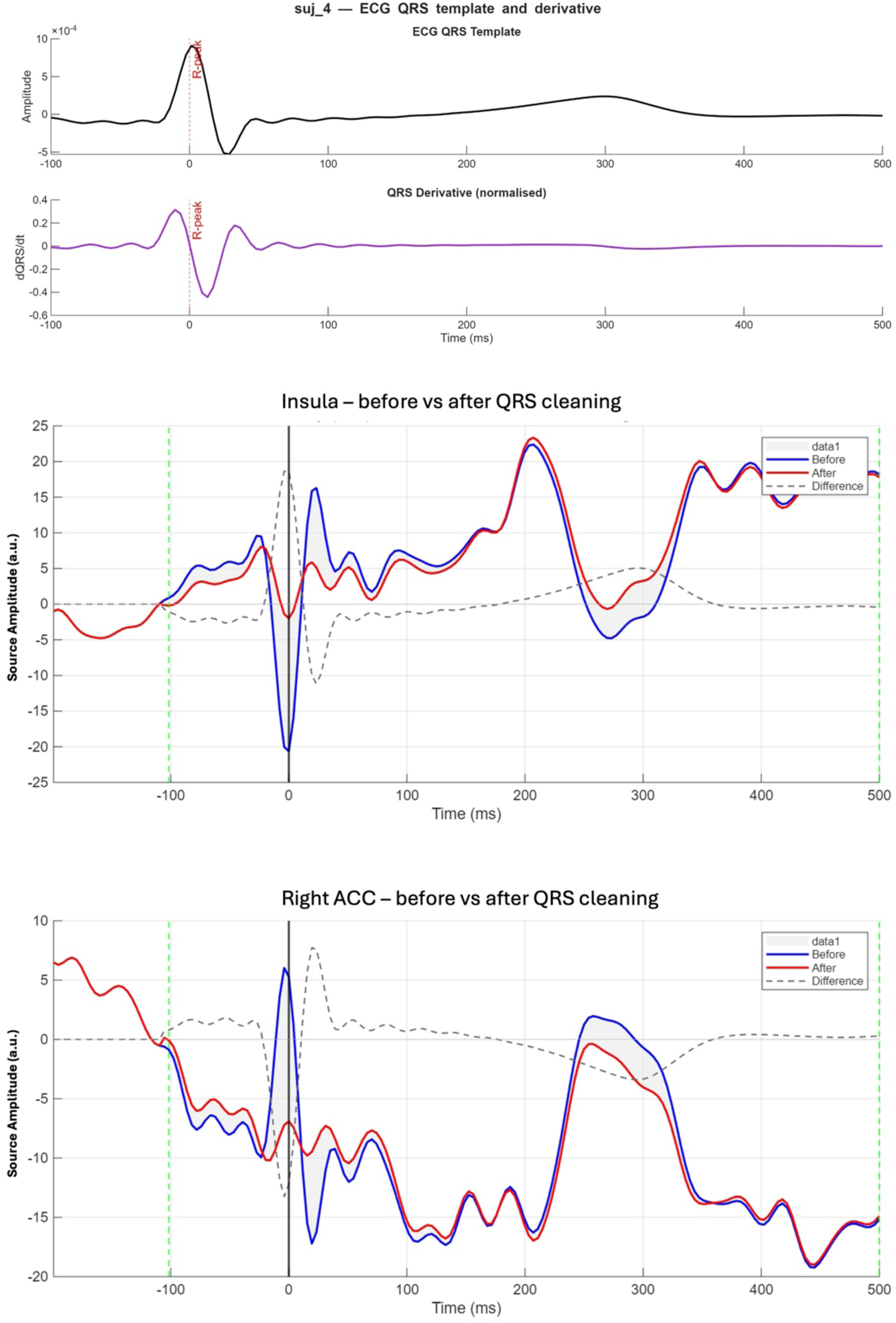
Example for one subject of the ECG template, its derivative and the source waveform for the right insula and right ACC VOIs before and after subspace correction.

As the output of LCMV beamforming yields source waveforms with an arbitrary sign, an iterative leave-one-out sign correction was applied separately per group and per ROI. For each participant in turn, the QRS-cleaned waveform was correlated with the leave-one-out grand average of the remaining participants; if the sign-flipped correlation exceeded the original, the waveform was negated. This was repeated until convergence (maximum 50 iterations).

#### 2.4.4. Signal-to-noise ratio estimation

To characterise the quality of the HEP signal in both sensor and source space, SNR was estimated for each simulated dataset and each subject from the empirical dataset using an alternating-sign noise floor method, which provides an empirical estimate of the residual noise in the averaged ERP.

Sensor space SNR – empirical data and simulations. For each simulation and subject, the real ERP was computed as the mean across all cardiac-locked epochs. To obtain a noise floor estimate, an alternating-sign ERP was computed such that odd-numbered trials were weighted +1 and even-numbered trials were weighted -1 before averaging. In sensor space, this would cancel out any consistent signal time-locked to the ECG, leaving a waveform dominated by residual noise. Channel B27 was identified as the scalp electrode with the strongest R-Ins projection via forward modelling of a noise-free simulation and was used for all subsequent sensor-space SNR calculations. Thus, sensor space SNR was computed on channel B27 as the ratio of the peak absolute amplitude of the real ERP to the RMS of the alternating-sign ERP, both evaluated within a 200-500 ms post R-peak window. This ensured that empirical and simulated data SNR values were computed on the same spatial and temporal basis, making them directly comparable.

Source space SNR – empirical data and simulations. Source-space SNR was computed from the beamformed counterparts of the same data used for sensor-space SNR estimation. For both empirical and simulated data, the alternating-sign averaged files were passed through the beamforming pipeline identically to the real averaged files. The noise floor in source space was defined as the RMS of the resulting beamformed alternating-sign waveform in the same 200-500 ms window post R-peak. Source-space SNR was computed as the ratio of the peak amplitude of the real beamformed ERP to this noise floor, expressed in dB.

Mapping real subjects onto the simulation SNR landscape. To contextualise each subject’s SNR relative to known waveform recovery performance from simulations, achieved sensor-space and source-space SNR from empirical data were plotted within a two-dimensional SNR landscape derived from the simulations. The centroid of each simulation condition in the SNR plane was defined as the mean achieved sensor-space and source space SNR across the 75 independent noise realisations, generating 20 control points for the interpolation. Using radial basis function interpolation with a thin-plate spline kernel, a smooth correlation surface was constructed by interpolating across the 20 simulated conditions. The colour of each point on the surface reflects the mean Pearson |*r*| between the beamformed and known R-Ins waveform observed at that simulation condition, providing a continuous estimate of expected waveform shape recovery fidelity across the SNR plane. Each empirical subject’s expected waveform correlation was estimated by evaluating this surface at their measured SNR coordinates. Subjects whose position corresponded to an expected |*r*|≥0.40 were designated as high-SNR subjects for whom beamforming was expected to yield reliable waveform shape recovery.

## 3. Results

### 3.1. Model1: single HEP source

#### 3.1.1. Waveform recovery

In the simplest case, a single HEP source in the R-Ins with no other competing source, LCMV beamforming recovered the known source with high fidelity across a wide range of SNR conditions (figure 7). In the absence of CA, mean waveform correlations exceeded r>0.98 (SD<0.002) at source level SNR ≥ -20dB and sensor level SNR ≥ -20dB, demonstrating near-perfect recovery under intermediate to good recording conditions. Performance degraded progressively as SNR decreased, with correlations falling to r ≍ 0.06 at the most challenging combination of source SNR -50dB and sensor SNR -30dB, suggesting a source space noise floor beyond which the beamformer cannot reliably recovery the target signal.

**Figure 7.**
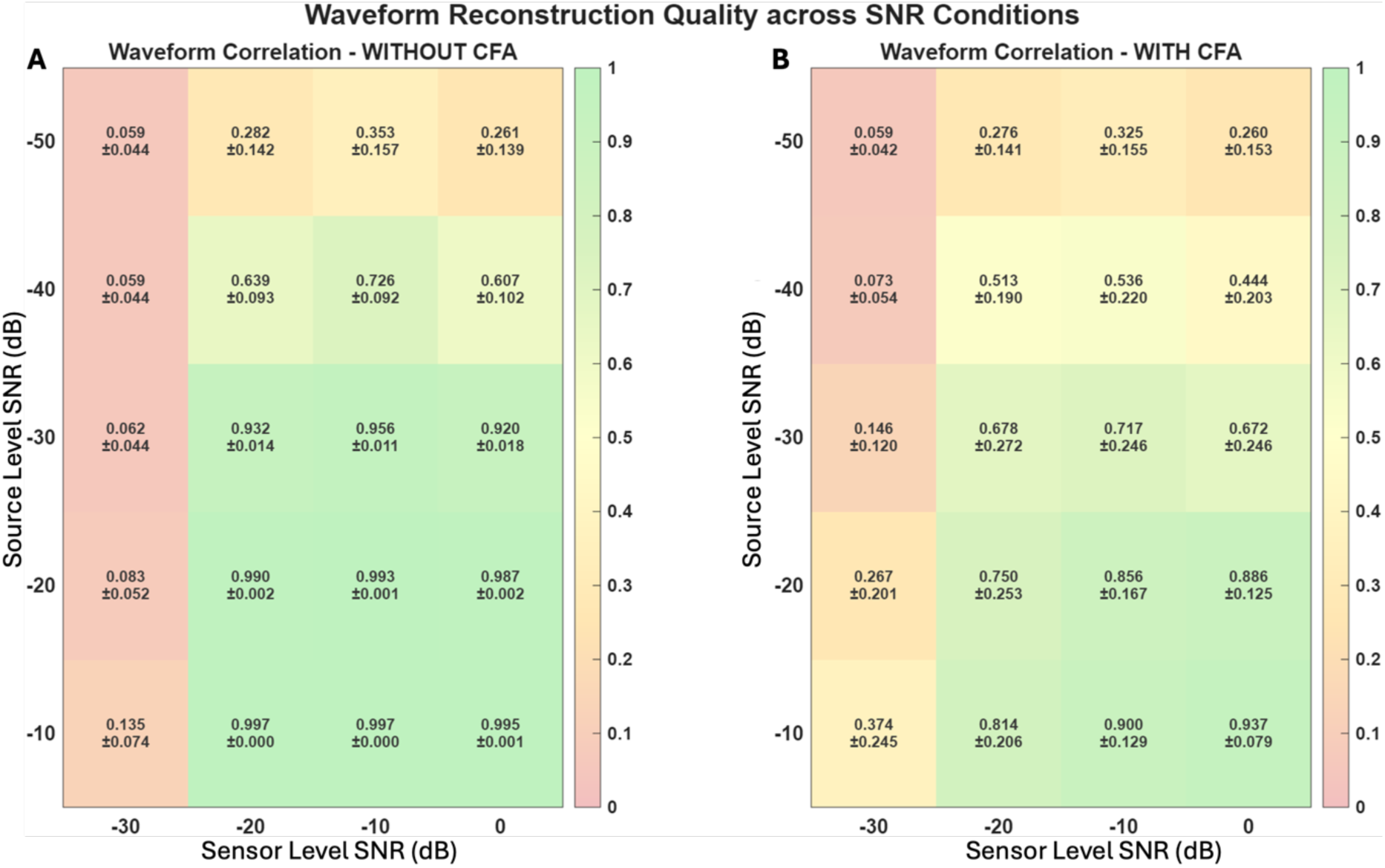
HEP waveform correlation to the known source without QRS cleaning across SNR conditions

Interestingly, the correlation between input waveform and beamformer output waveform were consistently lower at the lowest sensor noise level (SNR 0 dB) compared to sensor noise levels of -10 dB and -20 dB. This result was independent of source levels SNRs. This counter-intuitive finding suggests that moderate sensor noise may paradoxically stabilise beamformer performance, possibly by improving the conditioning of the sensor covariance matrix (Woolrich et al., 2011).

In the presence of CA (panel B), the broad SNR-dependent pattern seen without CA, was preserved but attenuated (figure 7). A strong correlation between input waveform and beamformer output was maintained (r=0.72-0.94) at source SNR≥-30dB and sensor SNR≥-20dB, although at lower SNRs this correlation was reduced (r<0.3 at sensor SNR <-20 dB). The correlation was higher in the presence of CA at source level SNR -30dB than without CA. This warrants caution in interpretation given the more striking finding: inter-individual variability increased substantially in the presence of CA (SD>0.2 in the presence of CA vs. SD <0.05 without CA across most conditions). This suggests that while beamforming is broadly resilient to CA, the degree to which individual CA characteristics affect waveform recovery is highly variable and not fully explained by SNR alone.

After QRS subspace cleaning (section 2.4.3) (figure 8), mean waveform correlations between input waveform and beamformer output were largely preserved and demonstrated modest improvements at intermediate to low SNR levels when CA was included in the simulation. Correlations marginally decreased at high SNR conditions. At source SNR -30 dB and sensor SNR -20dB, correlation improved from r = 0.678±0.272 to r = 0.743±0.217 after cleaning. At the highest SNR conditions, source SNR -10dB and sensor SNR 0dB, cleaning introduced a small decrease in correlation to the known input source (r = 0.901±0.122 vs r = 0.937±0.079). This suggests that CA subspace projection removes a marginal amount of signal alongside artefact at high SNR. QRS cleaning also reduced inter-individual variability at intermediate SNR levels. Overall, QRS cleaning demonstrated a modest but consistent benefit in simulated conditions where CA was most significant, without substantially altering the SNR-dependent performance pattern established in the original, non-QRS cleaned results.

**Figure 8.**
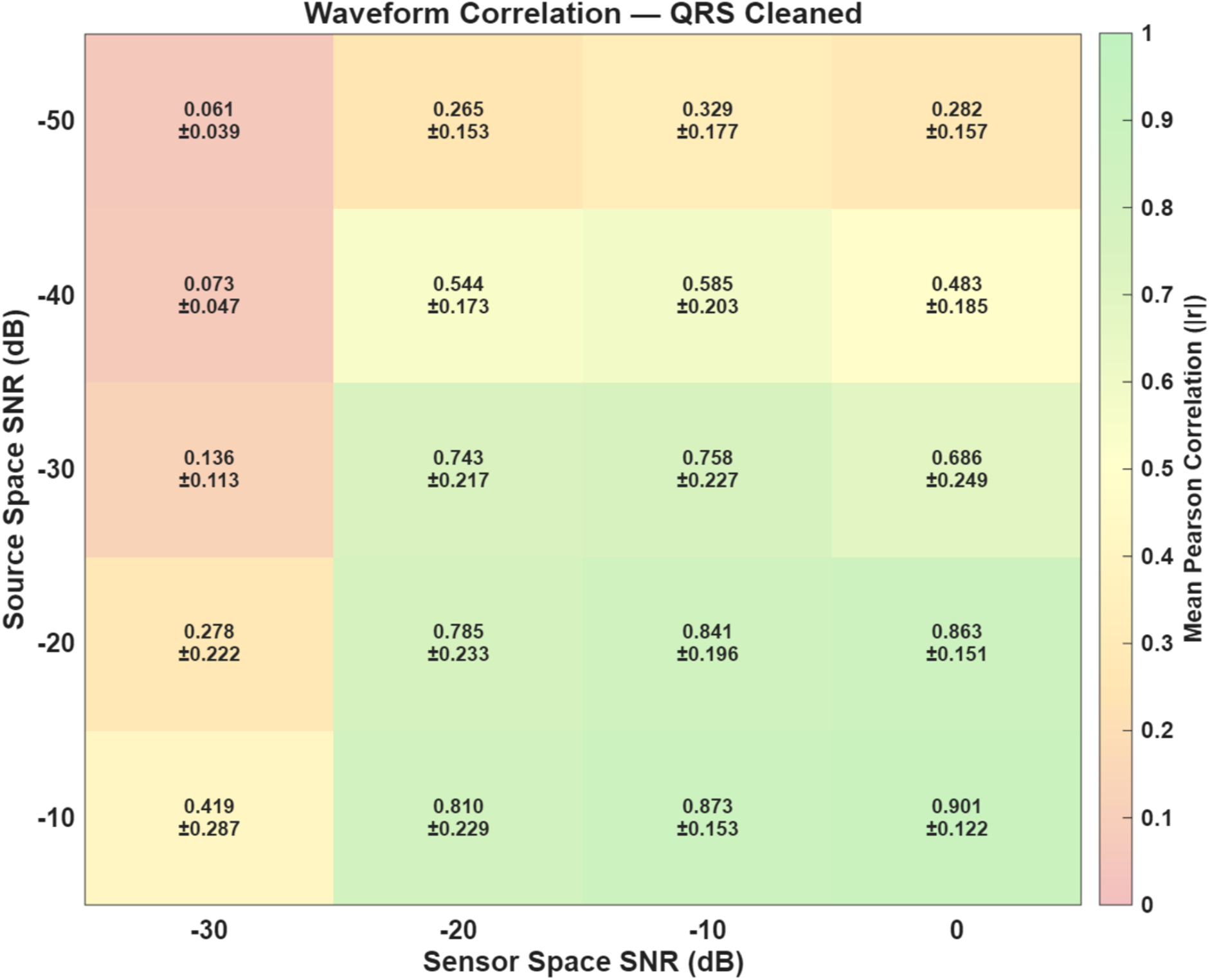
HEP waveform correlation to the known source after QRS cleaning

#### 3.1.2. Source localisation

In the absence of CA, insula source localisation error was negligible to minimal across most conditions up to source SNR -40dB (0-4.0mm). Localisation errors remained at ≤ 8.2mm at source SNR -50dB and sensor SNR -20dB (figure 9). In the presence of a single source, beamforming retrieves the known source within an anatomically meaningful distance of the true dipole across a wide range of SNR conditions. Robust performance was also found in the presence of CA, maintaining spatial source accuracy (0-5.7mm) at source SNR≥-30dB and sensor SNR≥-20dB. Localisation error increased at low SNR, particularly at −30 dB sensor SNR where errors of 16.2 mm were observed for both −40 and −50 dB source SNR. Localisation failure, defined as peak voxel distance exceeding 20 mm from the known source, occurred exclusively at sensor SNR -30 dB regardless of source SNR or CA.

**Figure 9.**
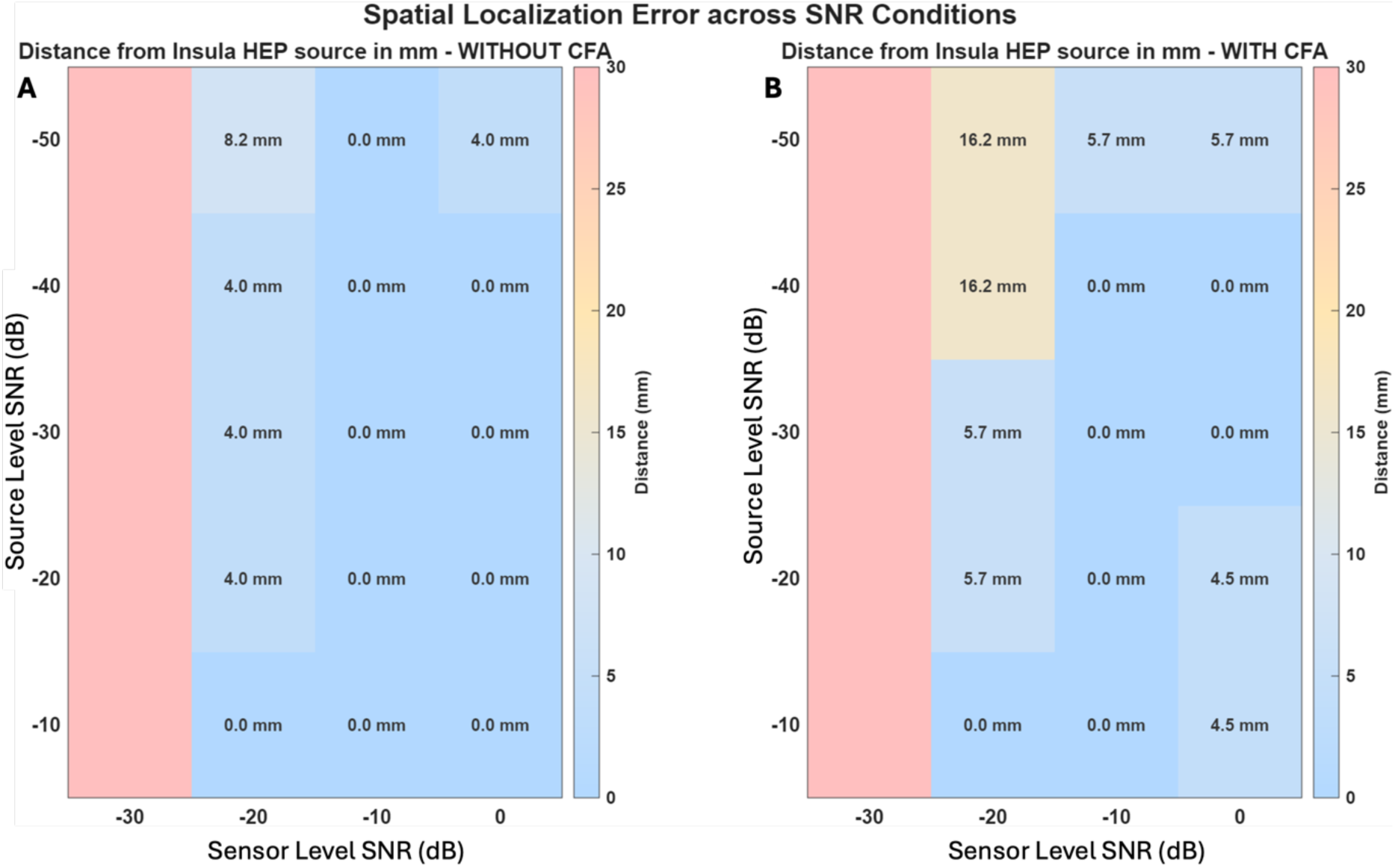
Spatial localisation error with respect to the known source location across SNR conditions

### 3.2. Model 2: two temporally distinct HEP sources

#### 3.2.1. Waveform recovery

Source waveform correlation results are in keeping overall with model 1 (figure 10). However, correlations were attenuated across all SNR levels. This is expected as the beamformer is now separating two competing sources.

**Figure 10.**
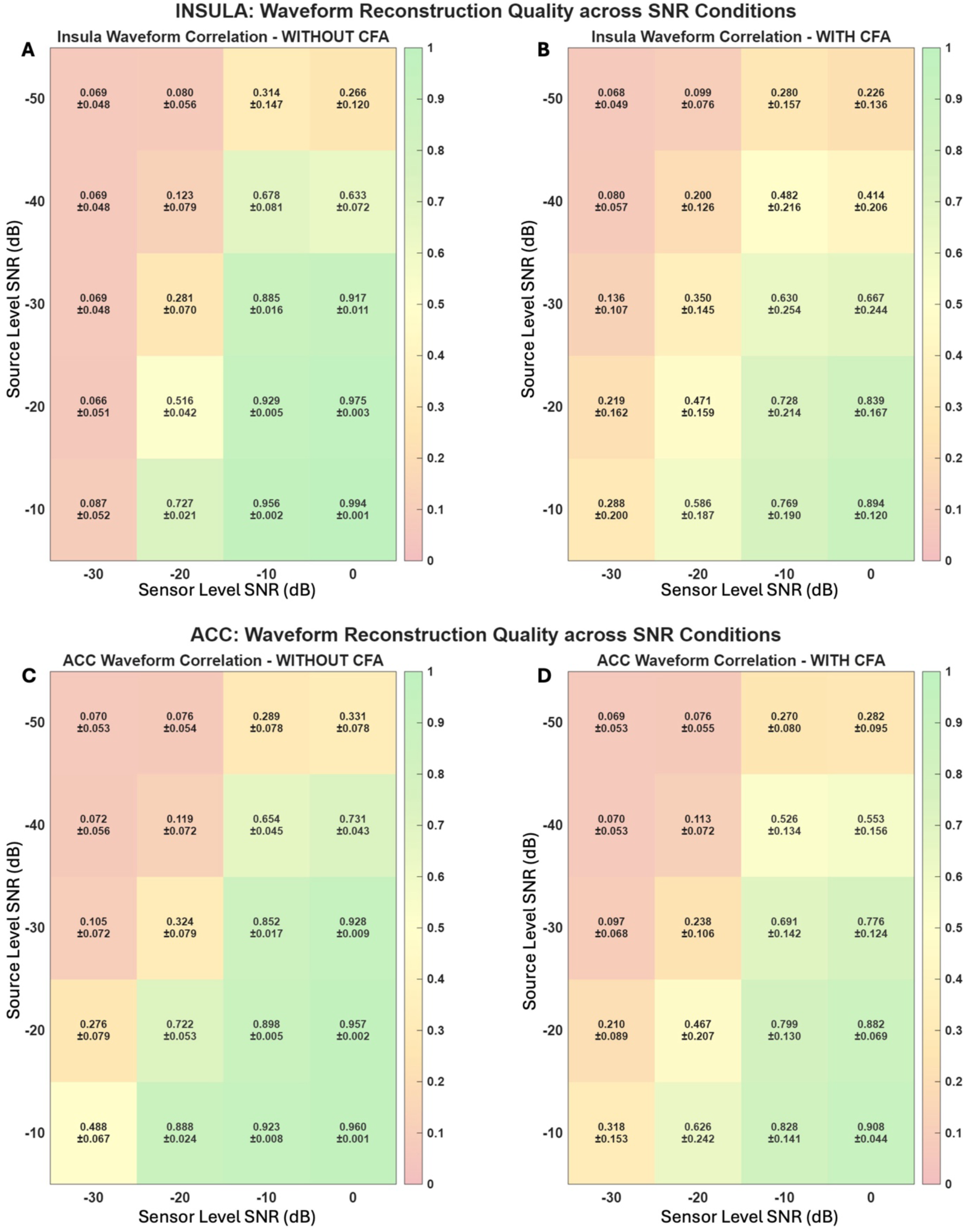
HEP waveform correlation to the known source without QRS cleaning across SNR conditions

Near-perfect correlation was maintained for the R-Ins source waveform recovery without CA (panel A) (r>0.885) up to source level SNR -30dB and sensor level SNR -10dB. Performance degraded more rapidly than in model 1 as SNR decreased, suggesting competition from the R-ACC source. For the right ACC source without CA (panel C), source correlation was higher at -20 dB sensor SNR than for the insula. This could suggest that beamforming is able to better locate the ACC source at low SNR or there is leakage of one into the other. The same pattern was observed for the ACC source in the presence of CA at high to intermediate SNRs. SD is low across all SNR combinations in the absence of CA.

Both the insula and the ACC source demonstrated high SDs in the presence of CA as in model 1. However, the beamforming performance for the ACC source in the presence of CA is notably better than for the insula source in the presence of CA – at sensor SNR 0dB and source SNR -10 dB r= 0.908 for ACC in the presence of CA vs r=0.894 for insula in the presence of CA. The source recovery is also more preserved from no CA to in the presence of CA for the ACC than for the insula – r=0.960 to r=0.908 vs r=0.994 to r=0.894 for insula. Waveform correlations for the insula in the presence of CA were high for source SNR ≥-20dB and sensor SNR≥-10dB (r>0.73). However, it degraded at a faster rate as SNR decreased than for model 1. This indicates competition with the ACC source. ACC source recovery without CA was comparable or superior to insula recovery across most SNR conditions with correlations reaching r=0.488 at source SNR -10dB and sensor SNR - 30dB vs r=0.087 for the insula. The temporal separability of the two sources conferred a clear localisation advantage for the ACC relative to the insula source under poor sensor noise conditions. Similarly to model 1, simulations in the presence of CA increased inter-individual variability with SD>0.1 for most conditions for the ACC and insula vs SD<0.08 without CA. Consistent with the spatial overlap between CA and insula, this pattern was more noticeable for the insula than for the ACC.

Following QRS cleaning (figure 11), waveform correlation in the presence of CA showed a consistent pattern of improvement for both the R-Ins and R-ACC sources across most SNR levels, though the effect differed between sources.

**Figure 11.**
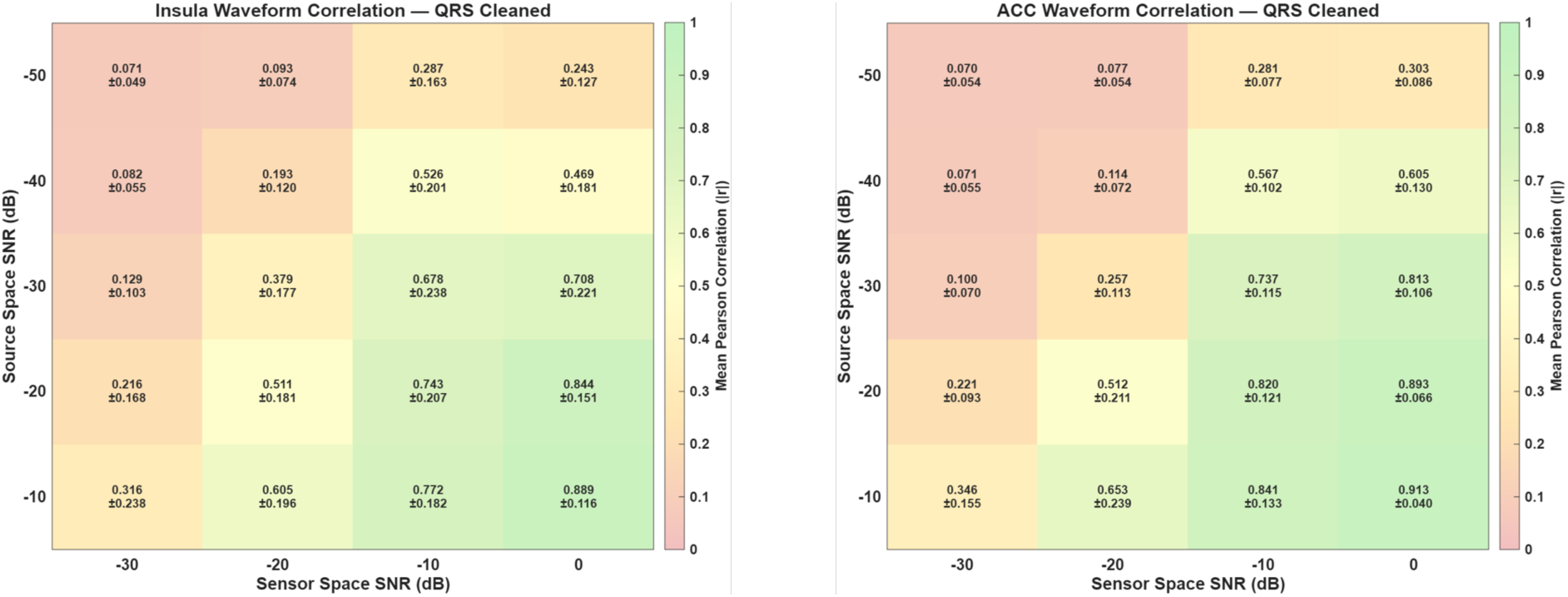
HEP waveform correlation to the known source after QRS cleaning

QRS cleaning generated meaningful improvements for the R-ins. At source SNR -20dB and sensor SNR 0dB, correlation improved from r = 0.839±0.167 to r = 0.844±0.151. At intermediate SNR (sensor SNR -10dB and source SNR -30dB), correlation increased after cleaning from r = 0.630±0.254 to r = 0.678±0.238. In line with model 1, subspace projection removed a small amount of signal at very high SNR. Inter-individual variability was reduced after cleaning across most intermediate conditions, thought high SDs persisted at sensor SNR -30dB.

For the R-ACC, QRS cleaning demonstrated larger and consistent improvements across all SNR levels compared to the R-Ins. At source SNR -20dB and sensor SNR 0dB, correlation improved from r = 0.882±0.069 to r = 0.893±0.066. At intermediate SNR (sensor SNR -10dB and source SNR -30dB), correlation increased after cleaning from r = 0.691±0.142 to r = 0.737±0.115. These results are consistent with the interpretation that CA overlaps more strongly with the insula spatial filter than the ACC filter, meaning that cleaning recovers more signal for the ACC, where the CA contribution to the source waveform was smaller to being with. As with the R-Ins, inter-individual variability remained high at sensor SNR -30dB regardless of cleaning, confirming that the worst sensor noise conditions remain the primary limiting factor.

#### 3.2.2. Spatial crosstalk between sources

Cross-correlation heatmaps (figure 12) were generated to investigate the ACC leakage into the insula source. This revealed significant leakage of R-ACC activity into the R-Ins spatial filter. Panel A shows that the R-Ins source waveform correlated strongly to the ACC known source, particularly at good source SNR combined with poor sensor SNR (r=0.790±0.034 at source SNR -10dB and sensor SNR -30dB; r=0.787±0.024 at source SNR -20dB and sensor SNR -20dB; r=0.659±0.038 at source SNR -30dB and sensor SNR -20dB). This suggests that high source amplitude does not protect against inter-source crosstalk when sensor noise is elevated, and that the two-source configuration introduces spatial aliasing not present in model 1. Importantly, the reverse leakage was substantially lower. Panels C and D for the right ACC source show weak cross-correlations (r≤0.372 without CA and r≤0.365 in the presence of CA) which seems to indicate the leakage is asymmetrical in the direction of the insula. This could be due to the near-collinear lead fields imposed by the parallel dipole configuration combined with the spatial geometry of these two regions. CA did not systematically worsen crosstalk but increased its variability as expected.

**Figure 12.**
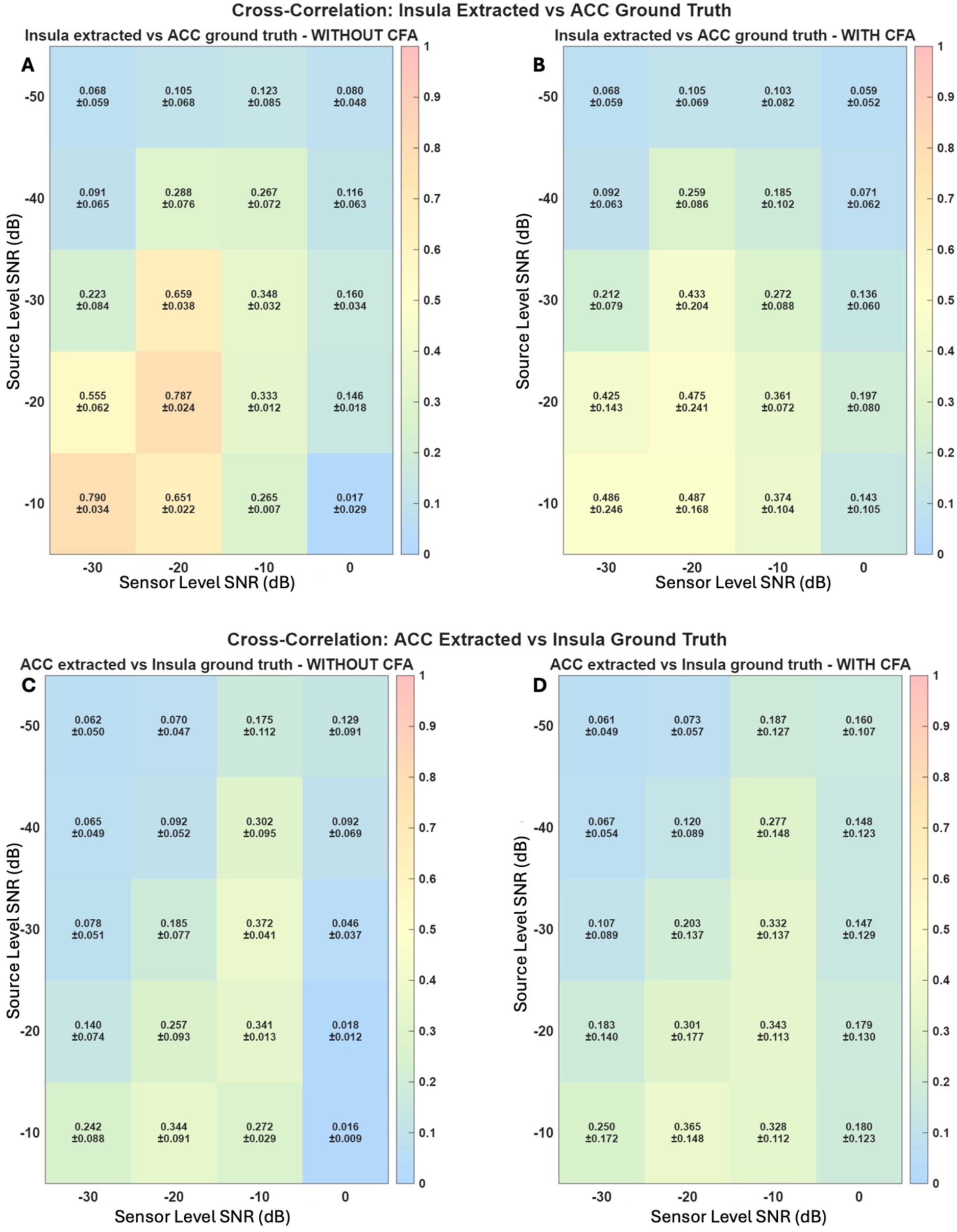
HEP waveform cross-correlation to the known source before QRS cleaning

#### 3.2.3. Source localisation

R-Ins localisation error was significantly increased compared to model 1 (figure 13). Localisation error exceeded 18.4 mm regardless of CA for sensor SNR ≤-10dB regardless of source SNR. At optimal SNR levels, localisation errors of 6 mm (sensor SNR 0dB and source SNR -10dB) were observed. This is most likely due to the crosstalk from the R-ACC source, pulling the beamformer peak away from the true R-Ins location (figure 12).

**Figure 13.**
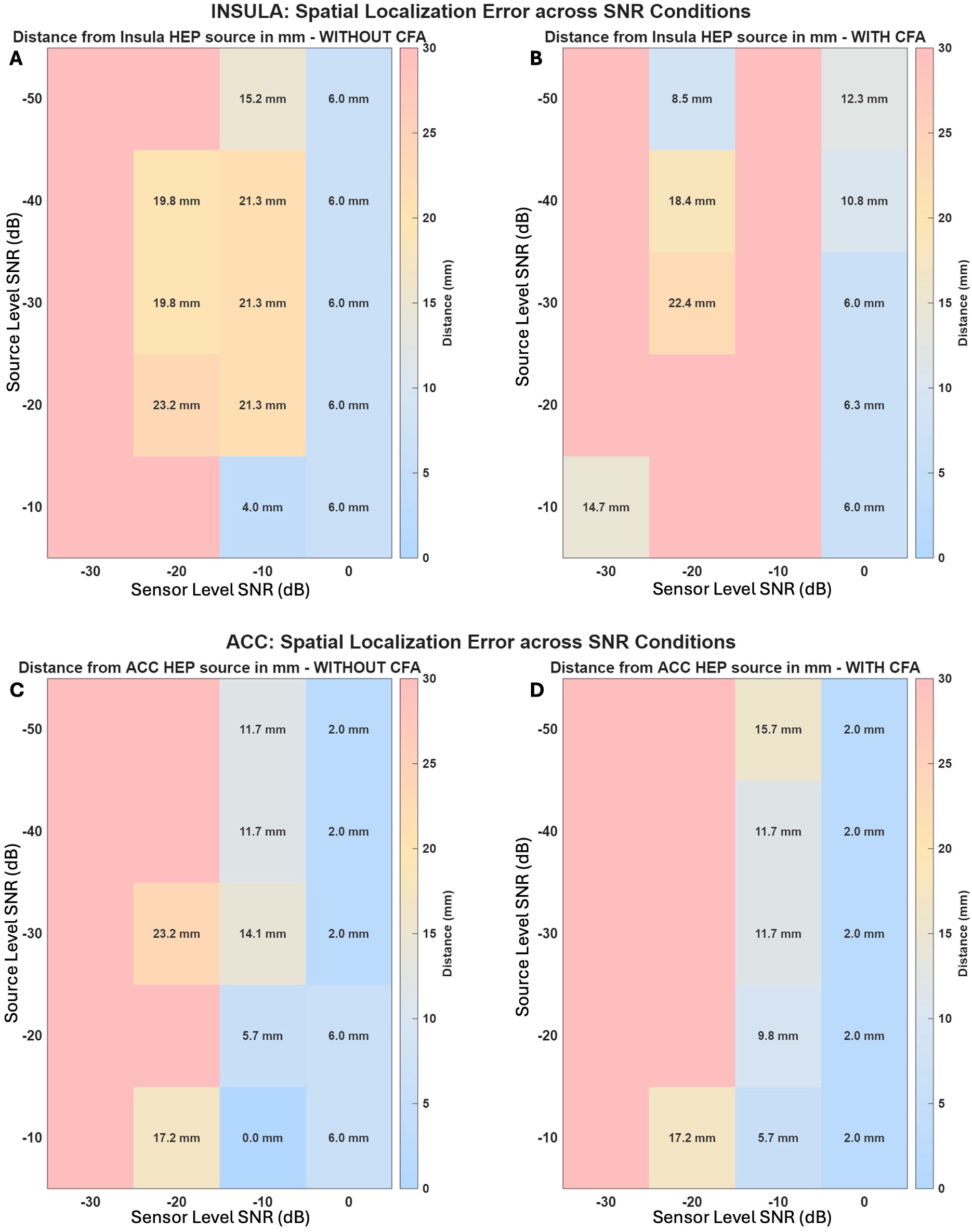
Spatial localisation error with respect to the known source location across SNR conditions

Conversely, R-ACC source localisation error without CA was lower than for the R-Ins (panel C): 5.7mm vs 21.3 mm respectively at source SNR -20dB and sensor SNR -10dB. CA (panel D) seemed to have minimal effect on R-ACC localisation accuracy and in some conditions, slightly improving it. This confirms the above speculations whereby the ACC is exerting a pull on the insula source resulting in bigger localisation errors for the insula source than for the R-ACC.

### 3.3. Model 3: two temporally overlapping HEP sources

#### 3.3.1. Waveform recovery

Waveform correlation results for both the insula and ACC sources were strikingly similar to those of model 2 across all SNR combinations (figure 14). Insula correlations with no CA ranged from negligible at the worst SNR conditions (r=0.059±0.045 at source SNR - 50dB and sensor SNR -30dB) to extremely strong (r=0.996±0.002 at source SNR -10dB and sensor SNR 0dB). This was similar for the ACC source with no CA: =0.067±0.045 at source SNR -50dB and sensor SNR -30dB and r=0.968±0.001 at source SNR -10dB and sensor SNR 0dB. In fact, correlation results were marginally stronger for model 3 than for model 2 for some SNR combinations: e.g. for ACC without CA and r=0.968±0.001 at source SNR -10dB and sensor SNR 0dB vs r=0.960±0.001 for model 2. This similarity to model 2 indicates that temporal overlap between HEP sources does not substantially impair the beamformer’s ability to recover individual source waveforms, consistent with the known reliance of the LCMV beamforming on spatial rather than temporal source structure.

**Figure 14.**
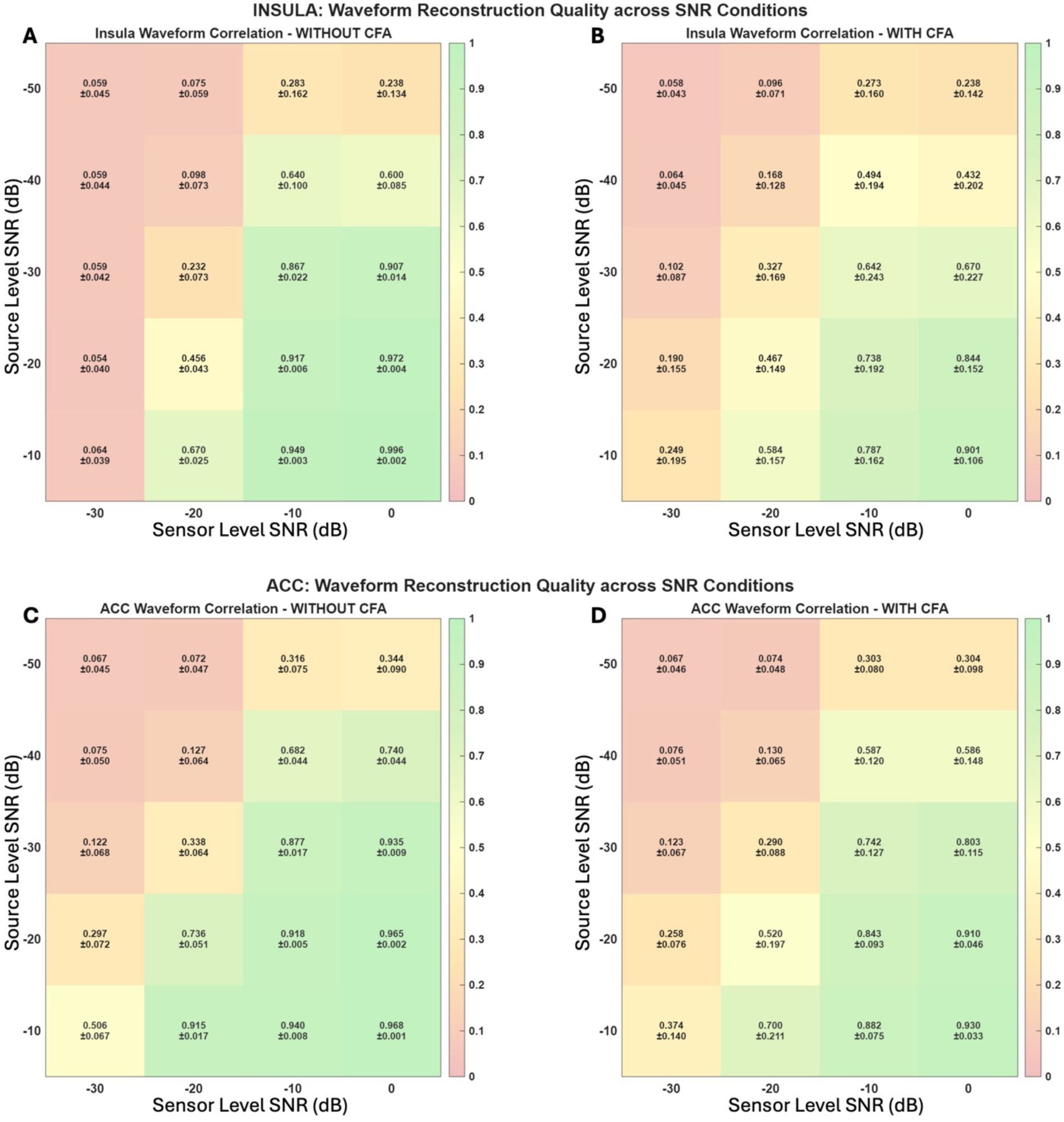
HEP waveform correlation to the known source without QRS cleaning across SNR conditions

As observed previously, correlations in the presence of CA showed substantial increased SDs for most SNR combinations: insula without CA and r=0.996±0.001 vs r=0.901±0.106 in the presence of CA at source SNR -10dB and sensor SNR 0dB. This confirms the inter-individual variability pattern observed in models 1 and 2. Panel D shows improved correlations in the presence of CA than to model 2 panel D in the presence of CA, suggesting that overlapping temporal sources could marginally aid the beamformer covariance estimation in some SNR conditions.

Following QRS cleaning in model 3 (figure 15), results mirrored observations in model 2, whereby the ACC source benefitted more consistently from cleaning than the insula source. Moderate improvements were observed for the R-Ins at intermediate SNR levels. At source SNR -30 dB and sensor SNR 0dB, correlation increased from r = 0.670±0.227 to r = 0.701±0.217. At source SNR -30dB and sensor SNR -10dB, correlation increased from r=0.642±0.243 pre-cleaning to r=0.689±0.208 post-cleaning. Results, broadly consistent with model 2, confirm that temporal overlap between sources does not meaningfully alter the effect of QRS cleaning on insula waveform recovery. Inter-individual variability after cleaning remained elevated at sensor SNR -30dB in line with models 1 and 2.

**Figure 15.**
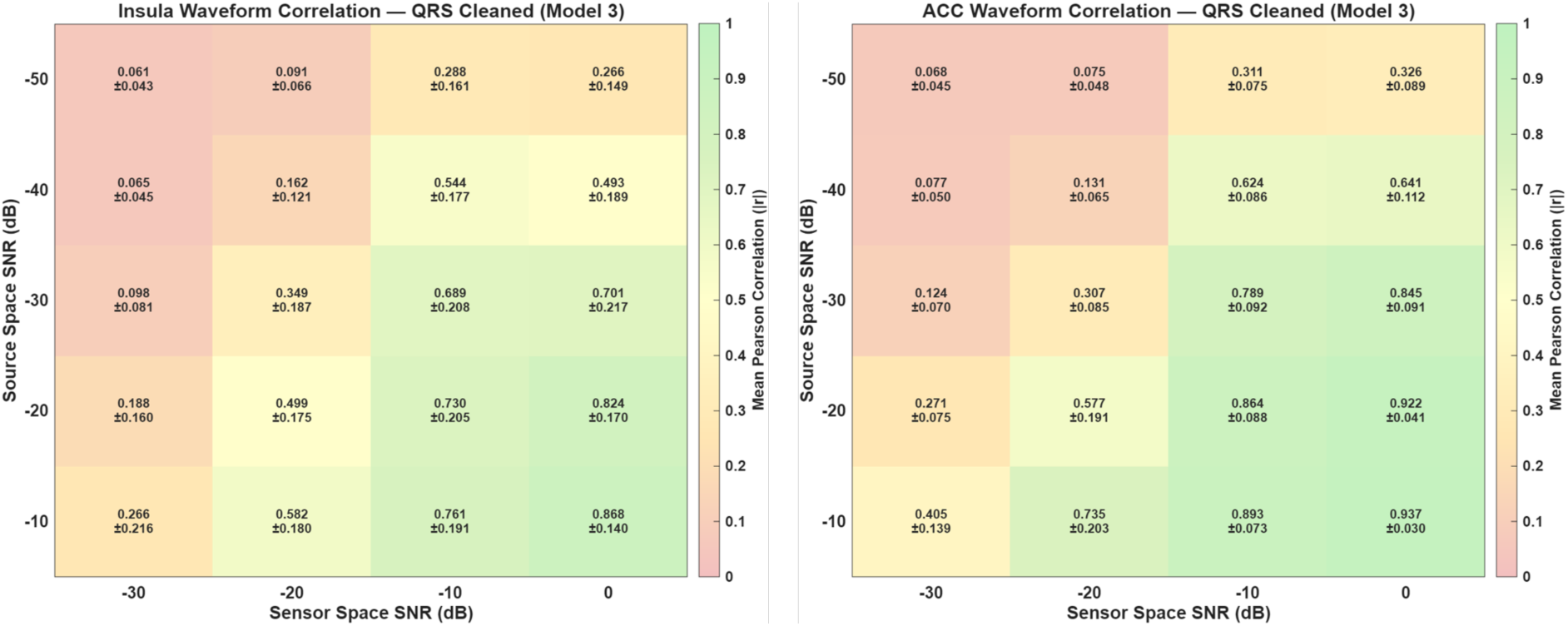
HEP waveform correlation to the known source after QRS cleaning

For the R-ACC, results were consistent with model 2, whereby consistent improvement in correlation was observed. At source SNR -30dB and sensor SNR 0dB, correlation improved from r = 0.803±0.115 to r = 0.845±0.091. Similarly, at source SNR -20dB and sensor SNR 0dB, correlation improved from r = 0.910±0.046 to r = 0.922±0.041. correlation increased at the highest SNR conditions as well: r = 0.930±0.033 to r = 0.937±0.030 at source SNR -10dB to sensor SNR 0 dB. Inter-individual variability was reduced across all conditions for the R-ACC. Correlations for the R-ACC exceed those of R-Ins, further extending the asymmetric pattern observed in model 2 and the interpretation that CA contamination preferentially affects the insula spatial filter relative to the ACC. In line with previous models, sensor SNR -30dB remained the primary performance boundary where QRS cleaning provided minimal benefit for either source.

#### 3.3.2. Spatial crosstalk between sources

Cross-correlation heatmaps (figure 16) indicated that simultaneous sources do not substantively increase spatial leakage of one source into the other. Insula-extracted source waveforms vs ACC known input source were nearly identical to results from model 2: r=0.803±0.033 at source SNR -10dB and sensor SNR -30dB; r=0.827±0.020 at source SNR -20dB and sensor SNR -20dB; r=0.684±0.034 at source SNR -30dB and sensor SNR -20dB). Similarly to model 2, the asymmetry was present whereby the insula spatial filter captures signal from the ACC source. This could further support the claim that spatial filter geometry has a greater impact on source leakage than the temporal relationship between sources. Temporal simultaneity of source does not meaningfully increase spatial crosstalk compared to temporally separated sources, which is an important positive finding for the validity of LCMV beamforming in the likely multi-source regime of HEP responses.

**Figure 16.**
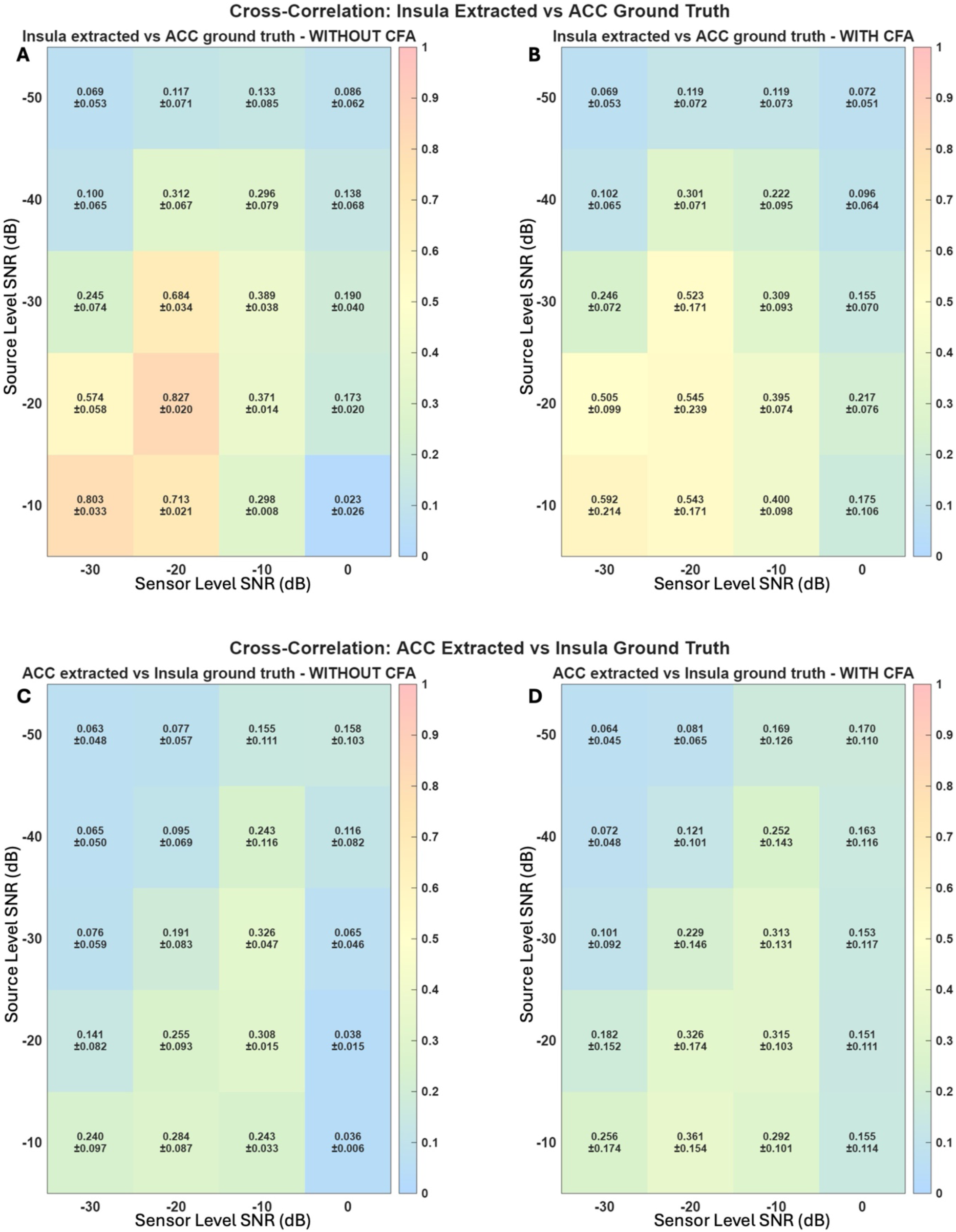
HEP waveform cross-correlation to the known source before QRS cleaning

#### 3.3.3. Source localisation

Temporal overlap between sources selectively degraded insula localisation relative to model 2 (figure 17). For the insula, with and without CA, beyond sensor SNR -10dB and source SNR -10dB, localisation error was above 22 mm. ACC localisation in model 3 was unchanged from model 2: 2.0 mm errors at 0dB sensor SNR across all source SNR levels and comparable partial recovery at -10dB sensor SNR. This dissociation, temporal overlap worsening R-Ins but not R-ACC localisation, further supports the interpretation that the insula filter is selectively sensitive to the covariance structure introduced by the concurrent ACC source. Comparatively, the R-ACC filter maintained spatial selectively regardless of the temporal relationship between sources.

**Figure 17.**
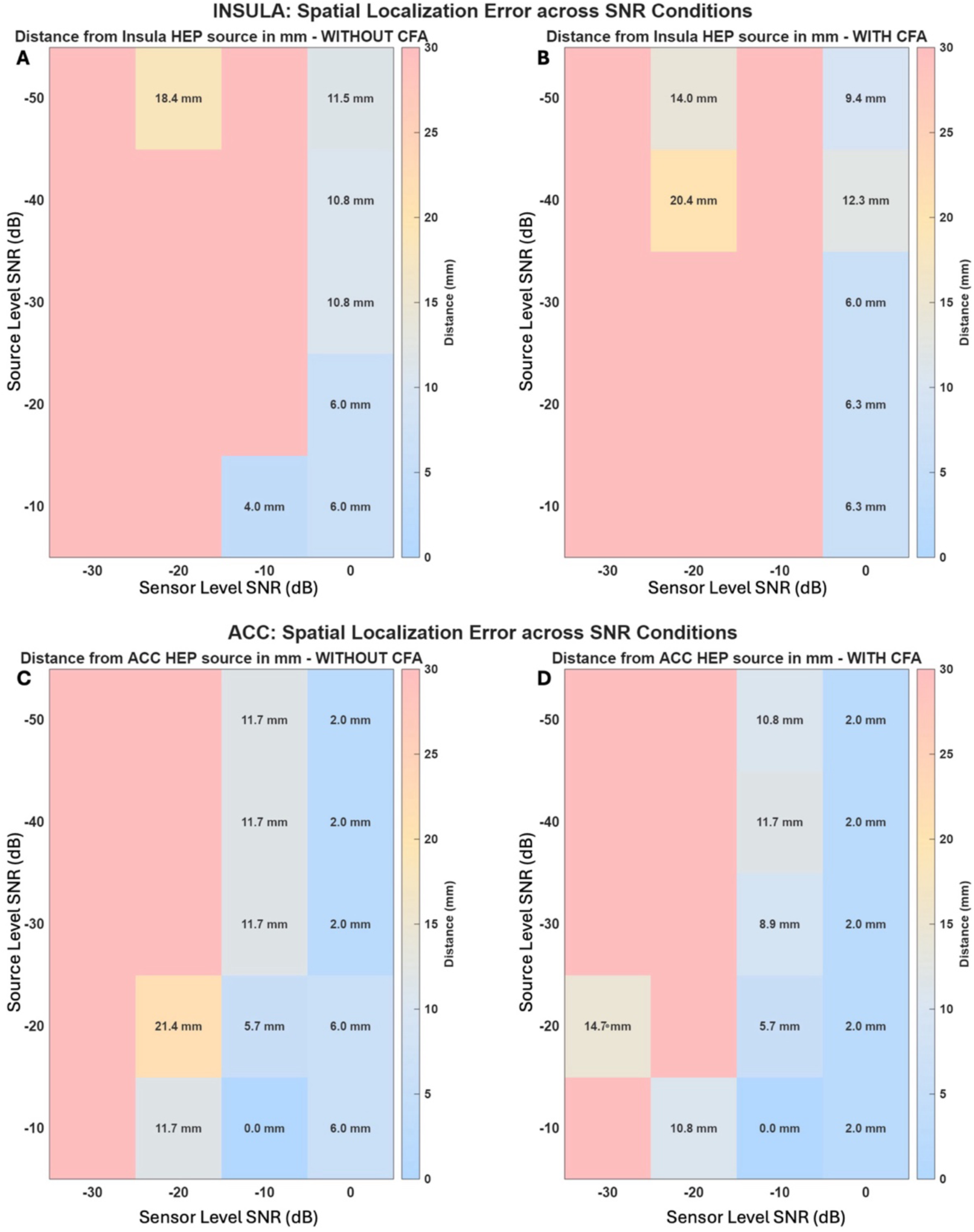
Spatial localisation error with respect to the know source location across SNR conditions

### 3.4. SNR quantification in empirical data and mapping onto the simulation landscape

Sensor-space and source-space SNR were estimated for all 106 individuals using the alternating-sign noise floor method described in 2.4.4. and plotted onto the simulation correlation landscape derived from model 1.

Source space SNR, computed from source extracted R-Ins time series, was broadly consistent across participants (mean 15.1±4.6 dB, median 15.3 dB, range 4.9 to 26.2 dB, IQR 12.3-18.6dB). Distribution was tight with 71% of subjects falling within a 10 to 20 dB band. This suggests beamforming is able to suppress noise in a relatively uniform manner. Sensor space SNR, estimated at channel B27, was substantially more variable (mean 15.7±6.9 dB, median 15.8 dB, range -0.6 to 31.9.2 dB, IQR 12.6 to 19.8 dB). Standard deviation was higher in sensor space compared to source space (6.9 vs 4.6 dB). The range of SNR was wider in sensor space than in source space with some subjects yielding a negative sensor-space SNR.

Mapping results onto the simulation landscape revealed that empirical data occupied a broad but coherent region of the source and sensor SNR plane (figure 18). Most subjects varied vertically across the plane reflecting sensor-space SNR variability. Fifty-eight subjects (∼ 55%) met the high-SNR criterion defined as reaching a correlation of |*r*|≥0.40, corresponding to moderate to high HEP recovery. The remaining 48 subjects (∼ 45%) clustered within the low SNR region of the source- and sensor-SNR plane, where simulations forecasted poor or unreliable HEP recovery. This simulation landscape provides a principled basis for interpreting the inter-individual variability in beamforming outcomes in the empirical data results that follow.

**Figure 18.**
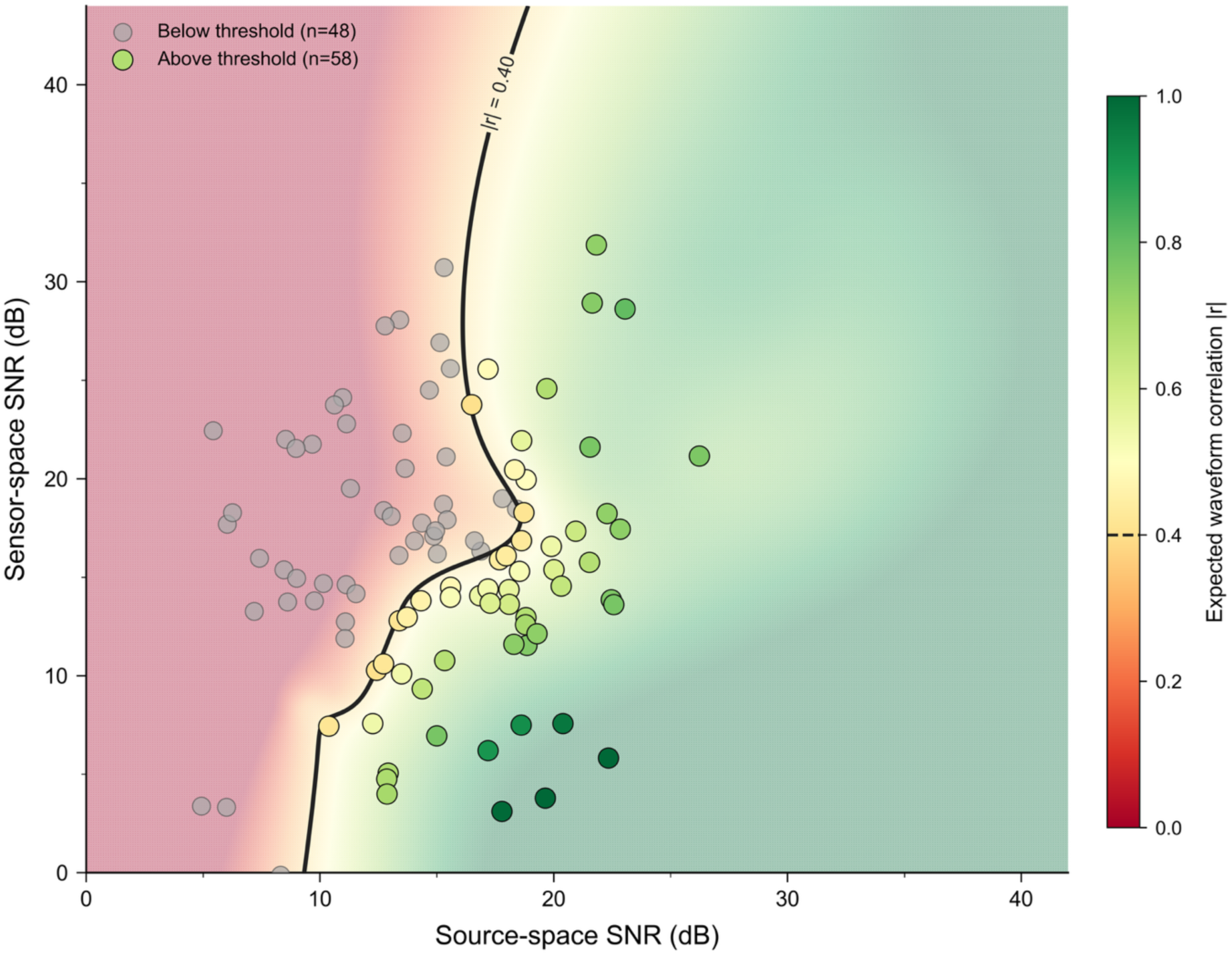
relationship between sensor-space and source-space SNR, across all subjects (n=106). Each point represents one subject, coloured by expected waveform correction |*r*| based on the simulation results from model 1. The black curve marks the correlation threshold, |*r*| = 0.4. Subjects (yellow and green points) falling to the right of the boundary (n=58), were deemed to have sufficient SNR in both domains to yield a reliable source-space waveform. Subjects (grey points) to the left of the black curve (n=48) did not meet this criterion and were excluded from subsequent source-space analyses. The simulated background gradient landscape reflects the expected |*r*| as a continuous function of both SNR dimensions.

### 3.5. HEP: pipeline validation and group comparisons

Results are presented in a 2 x 2 x 2 factorial design varying three factors: spatial projection (sensor space vs source space LCMV beamforming), cardiac signal null space projection (i.e. no QRS cleaning / with QRS cleaning), and subject inclusion (all subjects vs excluding poor SNR subjects if |*r*| < 0.4). This generated 8 distinct pipelines, applied to two separate clinical datasets as presented in section 2.4.1. (ANX vs NANX and HTN vs NHTN) (table 1). Similarly to the models 2 and 3, source extraction was performed in the R-Ins and R-ACC. Pooled analysis across all clinical samples was also performed using a one-sample t-test. For all group comparisons, Welch’s independent-samples t-test was the primary statistic. Mann-Whitney U tests were computed as a non-parametric verification. Results were consistent throughout and are not reported in detail. Cluster-corrected permutation tests (5,000 permutations, cluster-forming threshold α=0.05) were performed at source level for each comparison. Sensor-space HEP was derived to confirm the presence of a detectable cardiac-locked cortical response and to establish a baseline for comparison with the source extracted waveforms.

**Table 1.** Factorial design (2 x 2 x 2) generating 8 distinct pipelines.

|  | Beamforming |  |  |  |
| --- | --- | --- | --- | --- |
|  | No (Sensor Space) |  | Yes (Source Space) |  |
|  | Cardiac Cleaning |  | Cardiac Cleaning |  |
|  | No | Yes | No | Yes |
| **All Subjects** | **1** | **2** | **3** | **4** |
| **Excluding Poor SNR Subjects** | **5** | **6** | **7** | **8** |

#### 3.5.1. Pooled results – presence of the HEP

The grand average HEP waveform across all subjects (figure 19), showed a typical HEP morphology (pipeline 1): a large deflection approximately 200 ms post R-peak with a large QRS complex at t=0ms. Topographic maps at successive latencies confirmed that the HEP had a frontal scalp distribution, consistent with previously reported HEP topographies. The waveform was highly consistent across subjects reflected by the tight SEM bands in the frontal ROI grand average (figure 20).

**Figure 19.**
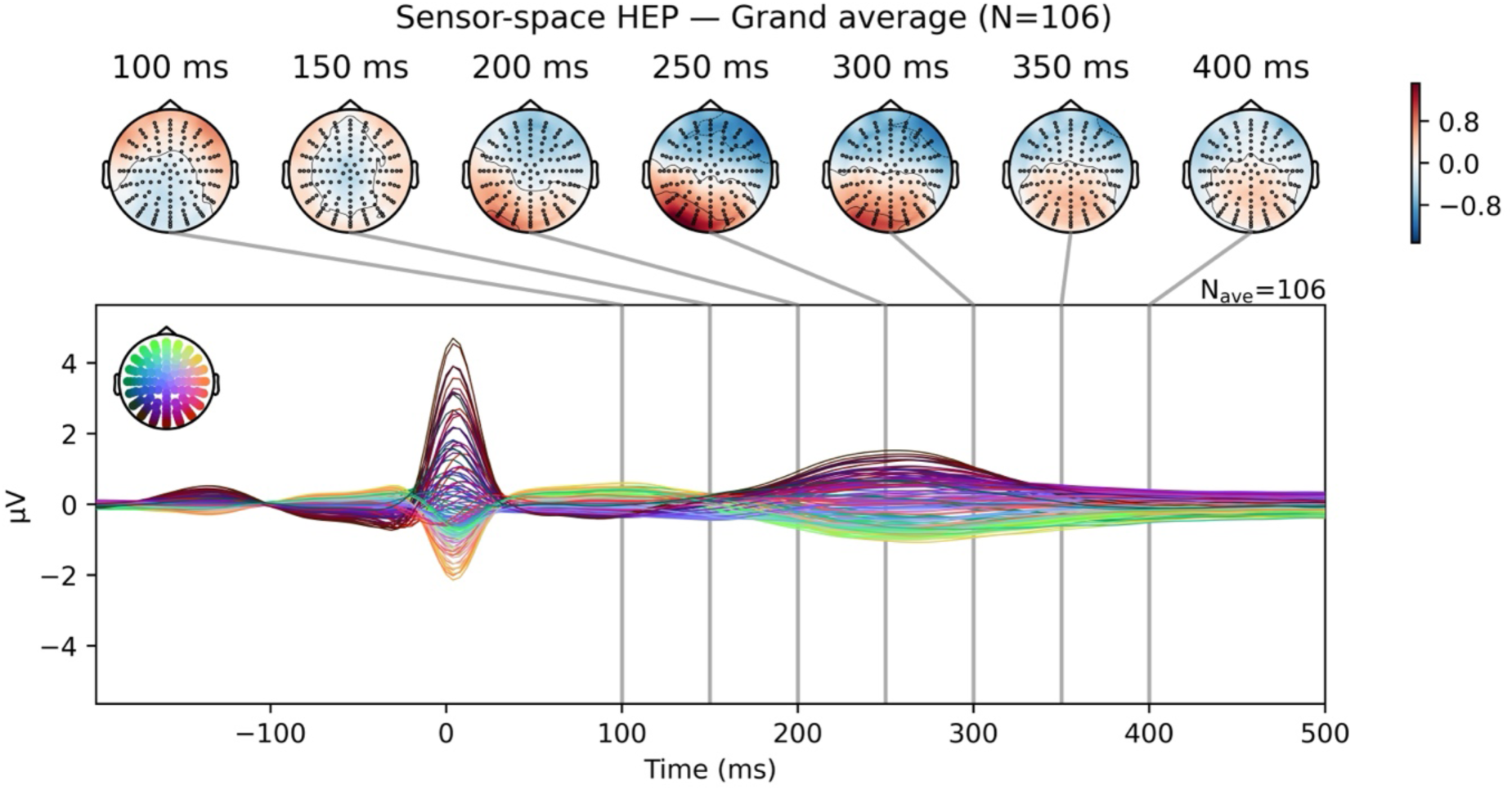
Sensor-space grand-average HEP waveform across all subjects (controls, ANX, HTN) N=106.

**Figure 20.**
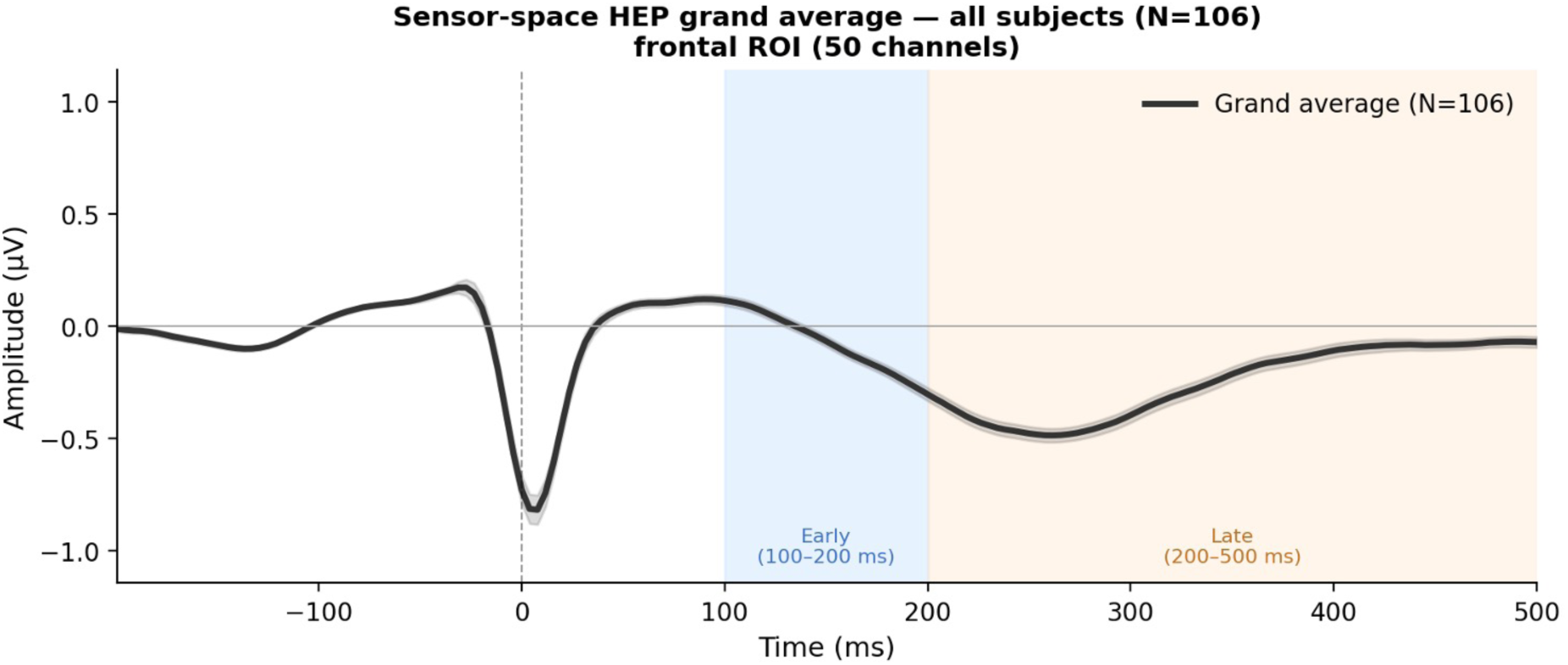
Grand-average BEP waveform across all controls, ANX and HTN subjects (N=106) computed as the mean across the frontal ROI (50 channels - C3,C4,C5,C6,C7,C9,C10,C13,C14,C15,C26,C27,C28,C31,C32,D3,D4,D5,D6,D7,C11, C12,C18,C19,C20,C21,C22,C23,C24,C25,D11,D12,D13,D14,D17,D18,D19,D20,D27,D 28,B17,B18,B19,B20,B21,B22,B23,B30,B1,B2). Shaded bands indicate ±1 SEM. Blue and orange shaded regions indicate the early (100-250 ms) and late (250-500 ms) HEP analysis window respectively.

To confirm that a reliable HEP can be detected using beamforming, one-sample t-tests were conducted over the frontal ROI amplitude pooled across four groups (ANX, NANX, HTN, NHTN) in sensor space as well as on the same pooled data in source space.

In sensor space, the HEP was highly significant across both time windows (early window 100-250 ms and late window 250-500 ms), both with and without QRS cleaning and for all subjects as well as when poor SNR subjects are excluded (p<0.001) (table 2, figure 21, figure 22). When all subjects were pooled together without QRS cleaning, the early HEP window yielded p<0.001, t(104) = -8.51, d = -0.83, and the late window p<0.001, t(104) = -8.08, d = -0.79. Following QRS cleaning, results remained significant (p<0.001) in both windows though HEP amplitudes were modestly attenuated (early window: -0.19 *μ*V vs - 0.15 *μ*V; late window: -0.21 *μ*V vs - 0.19 *μ*V). Results demonstrate a similar pattern when poor SNR subjects were excluded with all windows showing significance (p<0.001, d ranging from -0.75 to -0.88), and HEP minimally attenuated post QRS cleaning (early window: -0.20 *μ*V vs -0.15 *μ*V; late window: -0.24 *μ*V vs -0.22 *μ*V).

**Figure 21.**
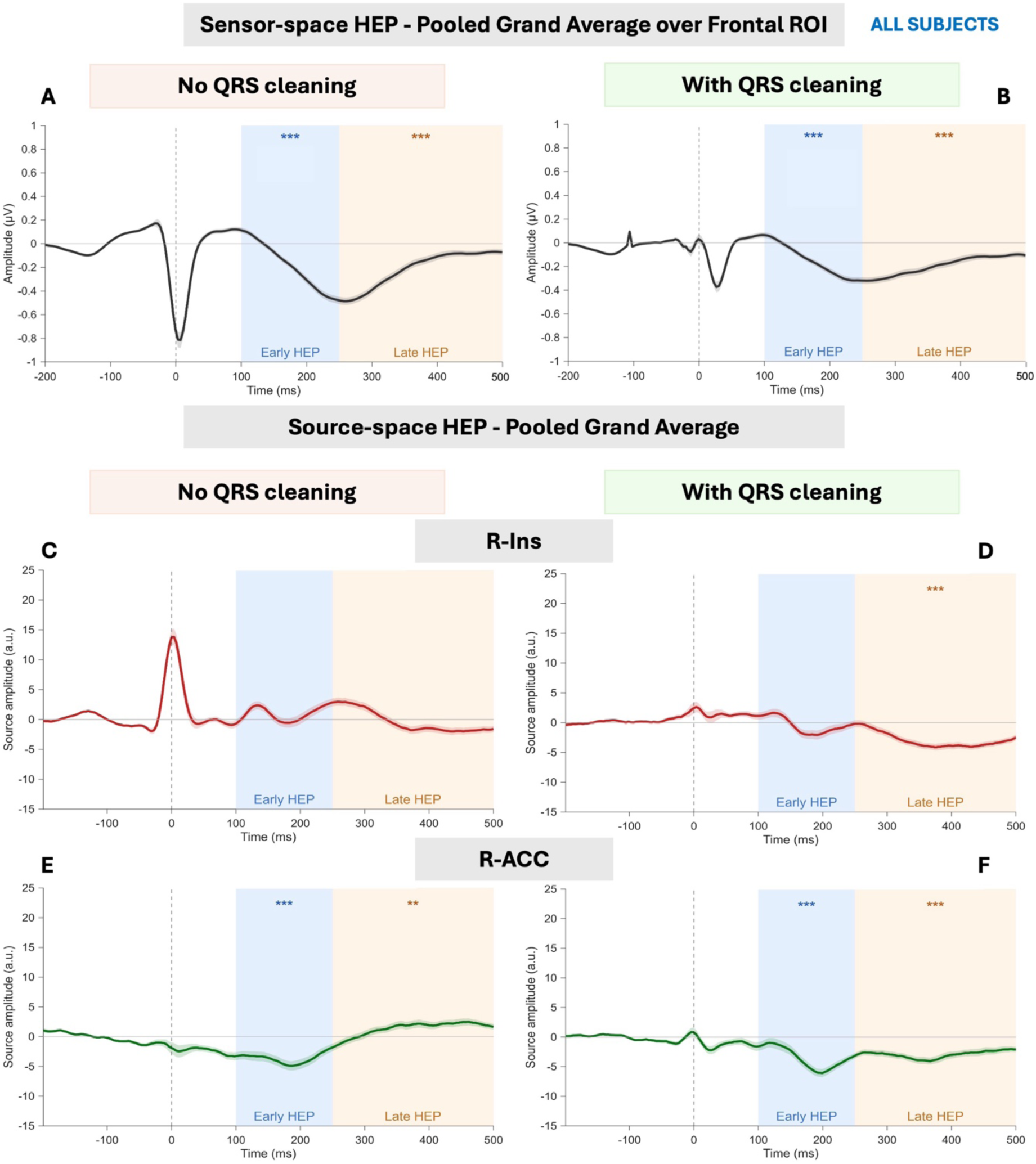
Pooled grand average HEP in sensor space and source space before and after QRS cleaning across all subjects.

**Figure 22.**
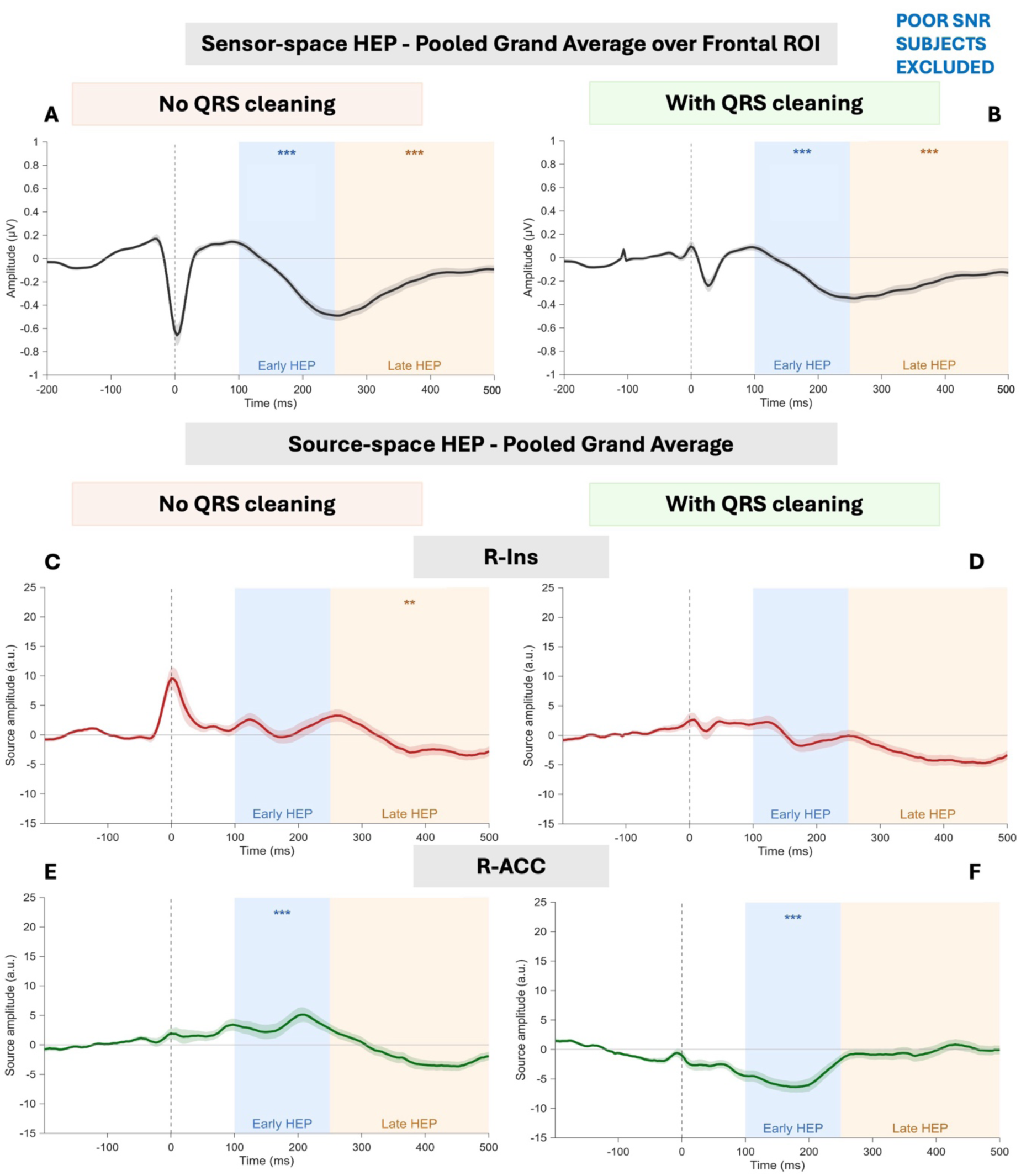
Pooled grand average HEP in sensor space and source space before and after QRS cleaning when poor SNR subjects are excluded.

**Table 2.** Pooled sensor-space HEP analysis: mean frontal ROI amplitude and one-sample t-test statistics across pipelines and subject cohorts.

| Window | Mean ($\mu V$) | SD | t(df) | p | Cohen's d |
| --- | --- | --- | --- | --- | --- |
| **No QRS cleaning - All subjects** |  |  |  |  |  |
| **Early** | -0.19 | 0.23 | -8.51 (104) | **<0.001** | -0.83 |
| **Late** | -0.21 | 0.27 | -8.08 (104) | **<0.001** | -0.79 |
| **QRS cleaned - All subjects** |  |  |  |  |  |
| **Early** | -0.15 | 0.21 | -7.48 (104) | **<0.001** | -0.73 |
| **Late** | -0.19 | 0.25 | -7.78 (104) | **<0.001** | -0.76 |
| **No QRS cleaning – Poor SNR subjects excluded** |  |  |  |  |  |
| **Early** | -0.20 | 0.22 | -6.70 (57) | **<0.001** | -0.88 |
| **Late** | -0.24 | 0.29 | -6.22 (57) | **<0.001** | -0.82 |
| **QRS cleaned - Poor SNR subjects excluded** |  |  |  |  |  |
| **Early** | -0.15 | 0.20 | -5.68 (57) | **<0.001** | -0.75 |
| **Late** | 0.22 | 0.27 | -6.22 (57) | **<0.001** | -0.82 |

In source space (table 3, figure 21, figure 22), the R-ACC showed a reliable HEP in both windows across most pipelines. In all subjects without QRS cleaning, the early window effect was highly significant (p<0.001, t(104) = -5.94, d = -0.58), as was the late window (p = 0.006, t(104) = 2.78, d = 0.27). Notably, after QRS cleaning, both windows remained significant (p<0.001), and effect sizes increased substantially in the late window (d = 0.27 vs d = -0.70). When poor SNR subjects were excluded without QRS cleaning, the R-ACC early window remained significant and was strengthened after QRS following (p=0.001, t(57) = 3.58, d = 0.47 vs p<0.001, t(57) = -5.15, d = -0.68). While the late R-ACC window was significant pre-QRS cleaning (p = 0.016), this effect disappeared after QRS cleaning (p = 0.58). This highlights that QRS cleaning eliminates spurious effects from CA contamination.

**Table 3.** Pooled source-space HEP analysis: mean frontal ROI amplitude and one-sample t-test statistics across pipelines and subject cohorts.

| Window | Mean (a.u.) | SD | t(df) | p | Cohen's d |
| --- | --- | --- | --- | --- | --- |
| **No QRS cleaning - All subjects** |  |  |  |  |  |
| **R-Ins Early** | 0.96 | 5.68 | 1.73 (104) | 0.087 | 0.17 |
| **R-Ins Late** | -0.34 | 5.08 | -0.72 (104) | 0.471 | -0.07 |
| **R-ACC Early** | -3.63 | 6.26 | -5.94 (104) | **<0.001** | -0.58 |
| **R-ACC Late** | 1.35 | 4.96 | 2.78 (104) | **0.006** | 0.27 |
| **QRS cleaned - All subjects** |  |  |  |  |  |
| **R-Ins Early** | -0.45 | 5.40 | -0.86 (104) | 0.392 | -0.08 |
| **R-Ins Late** | -2.96 | 4.08 | -7.44 (104) | **<0.001** | -0.73 |
| **R-ACC Early** | -3.51 | 5.99 | -6.01 (104) | **<0.001** | -0.59 |
| **R-ACC Late** | -2.93 | 4.21 | -7.14 (104) | **<0.001** | -0.70 |
| **No QRS cleaning – Poor SNR subjects excluded** |  |  |  |  |  |
| **R-Ins Early** | 1.29 | 6.57 | 1.49 (57) | 0.141 | 0.20 |
| **R-Ins Late** | -1.18 | 5.76 | -1.55 (57) | **0.045** | -0.20 |
| **R-ACC Early** | 3.38 | 7.19 | 3.58 (57) | **0.001** | 0.47 |
| **R-ACC Late** | 1.66 | 5.08 | -2.48 (57) | **0.016** | -0.33 |
| **QRS cleaned - Poor SNR subjects excluded** |  |  |  |  |  |
| **R-Ins Early** | -0.05 | 6.39 | -0.06 (57) | 0.95 | -0.01 |
| **R-Ins Late** | -3.25 | 4.80 | -5.15 (57) | **<0.001** | -0.68 |
| **R-ACC Early** | -5.04 | 5.70 | -6.74 (57) | **<0.001** | -0.89 |
| **R-ACC Late** | -0.39 | 5.36 | -0.56 (57) | 0.58 | -0.07 |

The R-Ins results revealed a distinct pattern. In the absence QRS cleaning, no significant pooled effect was observed in either window (early: p = 0.087, t(104) = 1.73, d = 0.17, late: p = 0.471, t(104) = -0.72, d = -0.07). Following QRS cleaning, a significant late HEP effect emerged (late: p<0.001, t(104) = -7.44, d = -0.73), while the early window remained non-significant (p = 0.392). This could indicate that QRS cleaning can reveal a genuine HEP signal masked by residual CA. When poor SNR subjects were excluded without QRS cleaning, a late-window insula effect in all pooled subjects was again significant (p = 0.045, t(57) = -1.55, d = -0.20) and was strengthened after QRS cleaning (p<0.001, t(57) = -5.15, d = -0.68). Pooled results indicate that CA can mask genuine HEP results in the R-Ins. QRS cleaning can unmask these signals rather than create them.

#### 3.5.2. Group comparisons

##### 3.5.2.1. Anxiety (ANX vs NANX)

###### Sensor Space

In sensor space, results for ANX vs NANX were not significant in any pipeline, window or cohort (all subjects and with poor SNR subjects excluded) (all p>0.13) (table 4, figure 23, figure 24). Without QRS cleaning in all subjects, the early HEP window showed p = 0.131, t(39.8) = -1.54, d = -0.42, and the late HEP window p = 0.311, t(33.6) = -1.03, d = -0.29. Results did not become meaningful with QRS cleaning (early: p = 0.340; late: p = 0.354). When poor SNR subjects were excluded, results were similarly non-significant. Cluster-corrected permutation tests revealed no significant results.

**Table 4.**
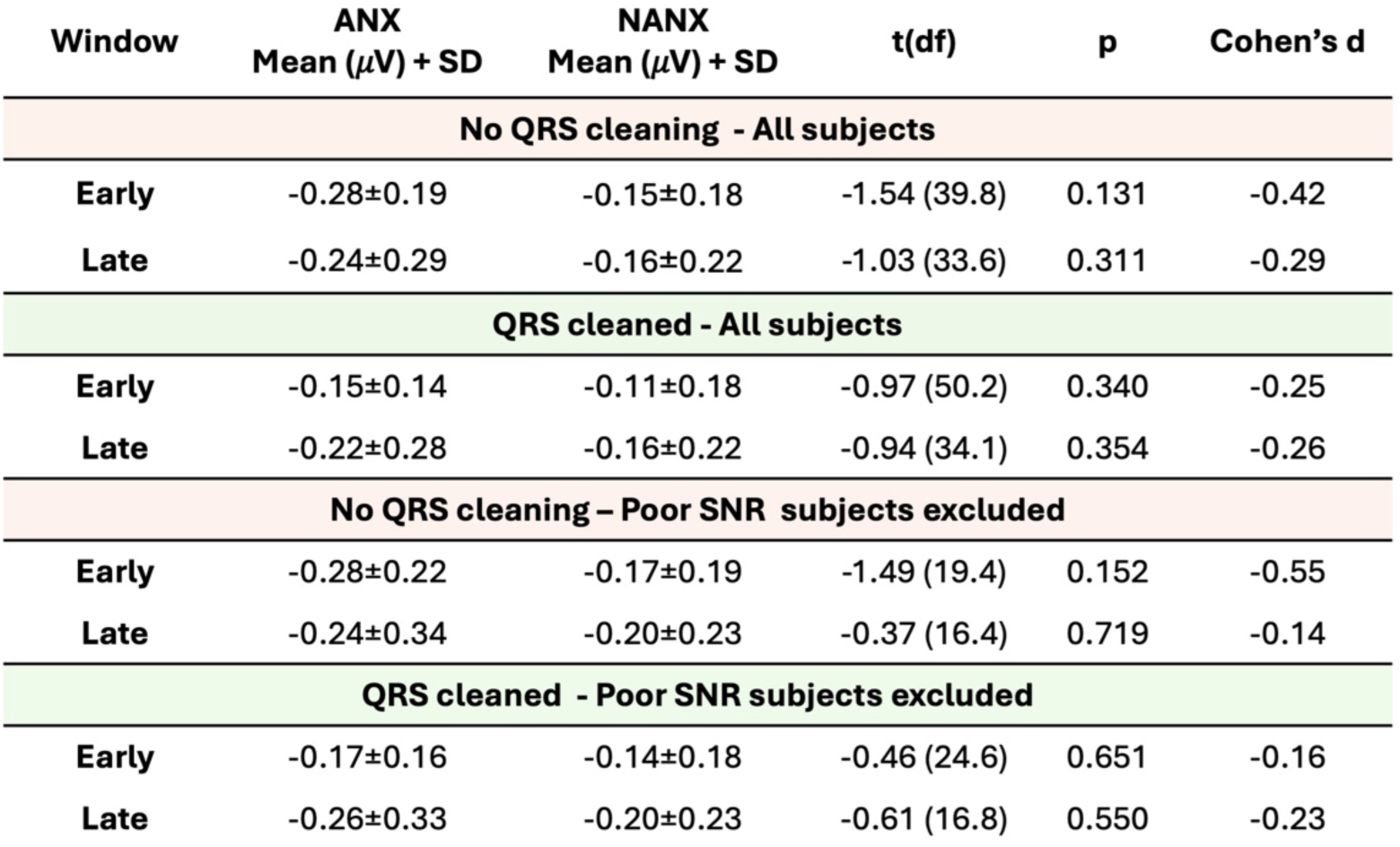
Sensor-space HEP analysis ANX vs NANX: mean frontal ROI amplitude and one-sample t-test statistics across pipelines and subject cohorts.

| Window | ANX<br>Mean ($\mu V$) + SD | NANX<br>Mean ($\mu V$) + SD | t(df) | p | Cohen's d |
| --- | --- | --- | --- | --- | --- |
| **No QRS cleaning - All subjects** |  |  |  |  |  |
| **Early** | -0.28±0.19 | -0.15±0.18 | -1.54 (39.8) | 0.131 | -0.42 |
| **Late** | -0.24±0.29 | -0.16±0.22 | -1.03 (33.6) | 0.311 | -0.29 |
| **QRS cleaned - All subjects** |  |  |  |  |  |
| **Early** | -0.15±0.14 | -0.11±0.18 | -0.97 (50.2) | 0.340 | -0.25 |
| **Late** | -0.22±0.28 | -0.16±0.22 | -0.94 (34.1) | 0.354 | -0.26 |
| **No QRS cleaning - Poor SNR subjects excluded** |  |  |  |  |  |
| **Early** | -0.28±0.22 | -0.17±0.19 | -1.49 (19.4) | 0.152 | -0.55 |
| **Late** | -0.24±0.34 | -0.20±0.23 | -0.37 (16.4) | 0.719 | -0.14 |
| **QRS cleaned - Poor SNR subjects excluded** |  |  |  |  |  |
| **Early** | -0.17±0.16 | -0.14±0.18 | -0.46 (24.6) | 0.651 | -0.16 |
| **Late** | -0.26±0.33 | -0.20±0.23 | -0.61 (16.8) | 0.550 | -0.23 |

**Figure 23.**
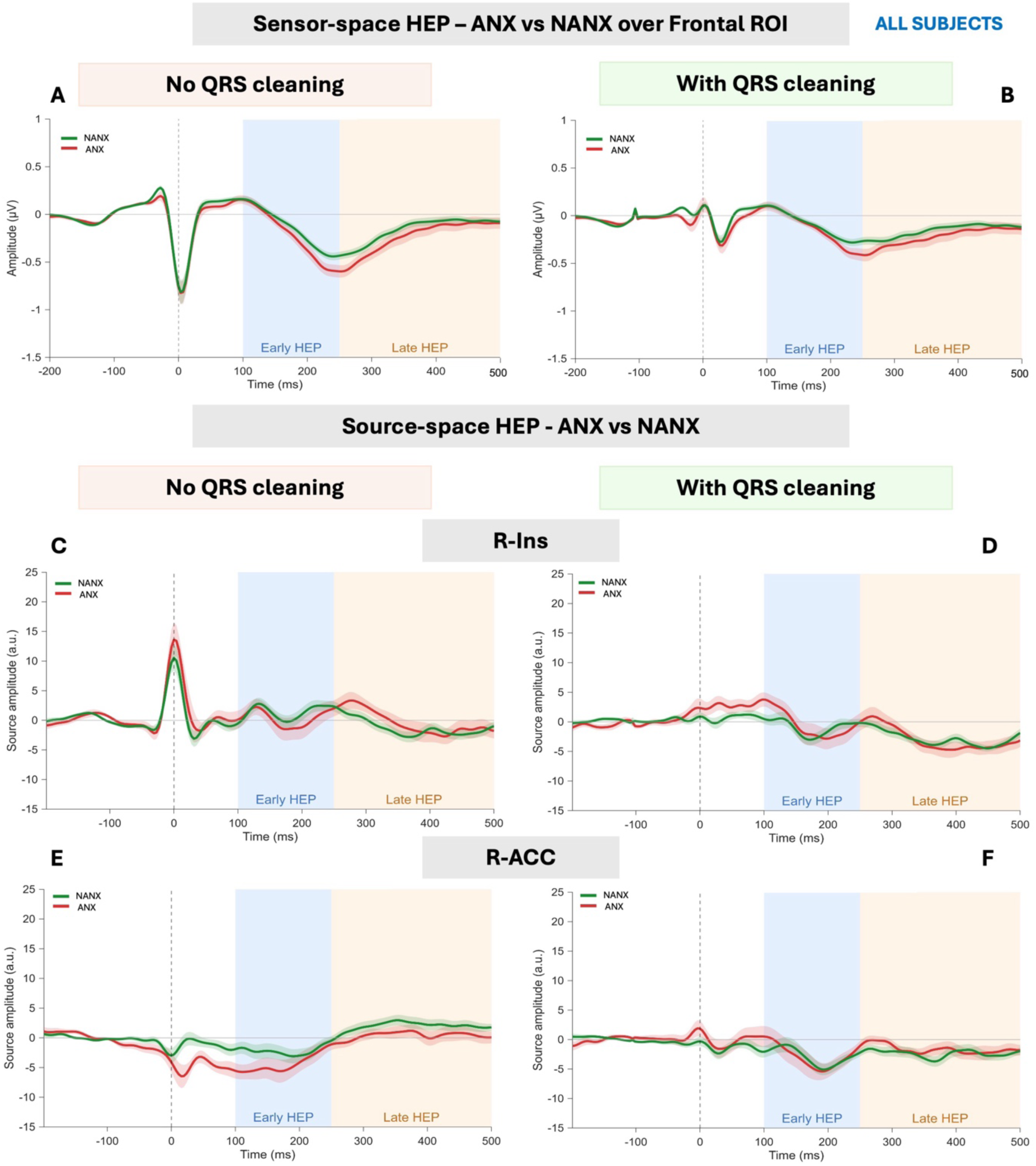
Grand average HEP in sensor space and source space before and after QRS cleaning across all subjects (ANX vs NANX).

**Figure 24.**
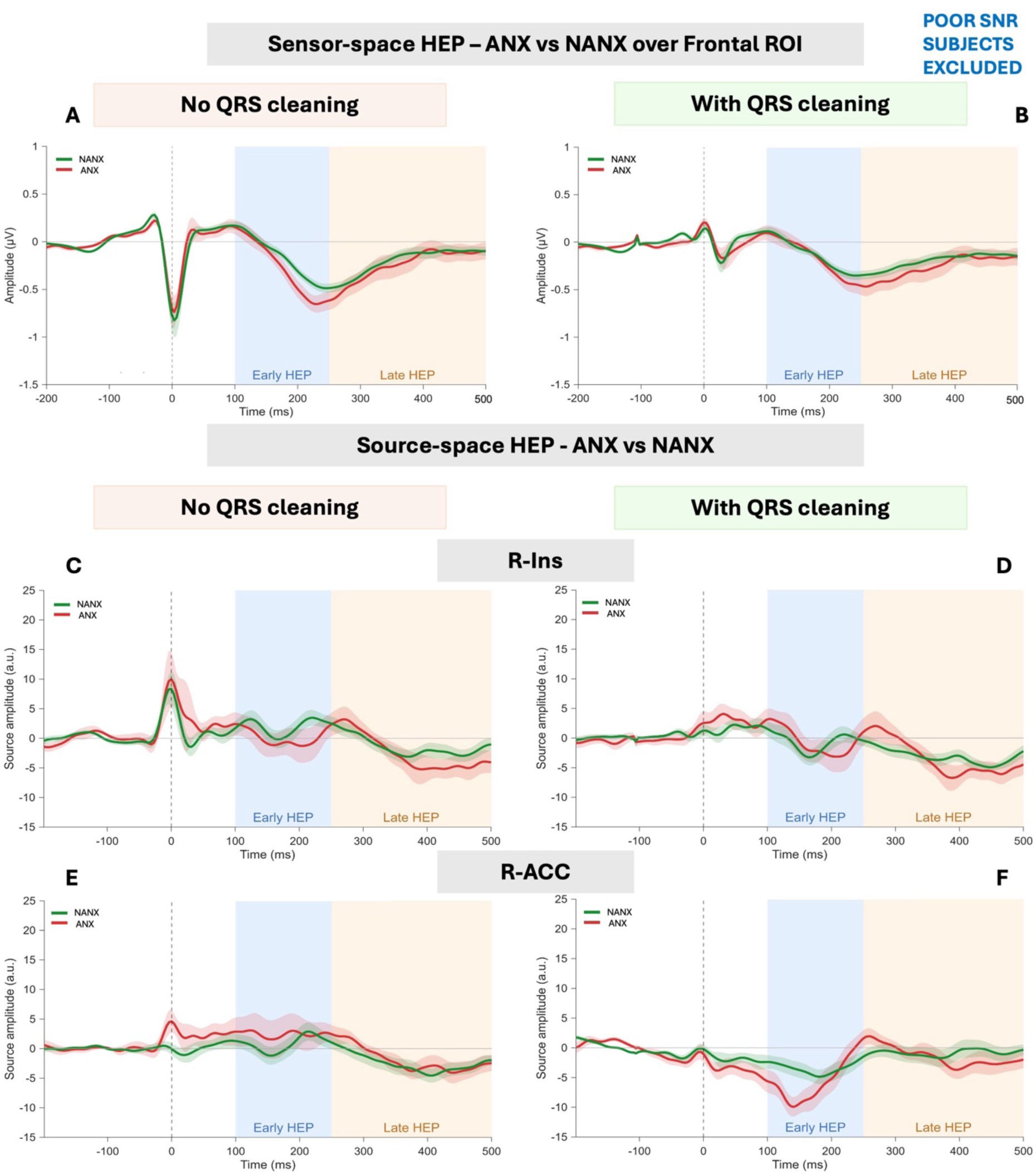
Grand average HEP in sensor space and source space before and after QRS cleaning when poor SNR subjects are excluded (ANX vs NANX).

###### Source Space

Source space results were consistent with sensor space results; no significant ANX vs NANX differences were observed in either VOI, time window, or pipeline (table 5, figure 23, figure 24). In the cohort with all subjects without QRS cleaning, the R-ACC window showed the largest effect, but this did not reach significance (p = 0.138, t(48.7) = -1.51, d = 0.40), and diminished after QRS cleaning (p = 0.445, t(50.3) = 0.77, d = 0.20). When poor SNR subjects were excluded, the R-ACC late window remained non-significant regardless of QRS cleaning (with QRS cleaning: p = 0.266, t(28.4) = 1.14, d = 0.39; without QRS cleaning: p = 0.200, t(27.7) = 1.31, d = 0.45). The insula showed no significant results in any pipeline. The findings suggest that LCMV beamforming does not artificially inflate effects, and that the null results in sensor space are preserved in source space.

**Table 5.** Source-space HEP analysis ANX vs NANX: mean frontal ROI amplitude and one-sample t-test statistics across pipelines and subject cohorts.

| Window | ANX<br>Mean (a.u.) + SD | NANX<br>Mean (a.u.) + SD | t(df) | p | Cohen's d |
| --- | --- | --- | --- | --- | --- |
| **No QRS cleaning - All subjects** |  |  |  |  |  |
| **R-Ins Early** | 0.33 ± 5.51 | 1.39 ± 4.22 | -0.77 (33.2) | 0.447 | -0.22 |
| **R-Ins Late** | -0.32 ± 4.99 | -1.24 ± 3.96 | 0.73 (34.1) | 0.473 | 0.20 |
| **R-ACC Early** | -4.39 ± 5.40 | -2.19 ± 5.20 | -1.52 (40.1) | 0.138 | -0.41 |
| **R-ACC Late** | 0.37 ± 3.54 | -1.95 ± 4.33 | -1.51 (48.7) | 0.138 | 0.40 |
| **QRS cleaned - All subjects** |  |  |  |  |  |
| **R-Ins Early** | -0.54 ± 5.69 | -0.93 ± 4.07 | 0.28 (31.5) | 0.784 | 0.08 |
| **R-Ins Late** | -2.92 ± 4.10 | -2.82 ± 3.16 | -0.10 (33.4) | 0.920 | -0.03 |
| **R-ACC Early** | -2.88 ± 6.12 | -2.88 ± 4.79 | 0.00 (33.8) | 1.000 | 0.00 |
| **R-ACC Late** | -1.63 ± 3.20 | -2.37 ± 4.09 | 0.77 (50.3) | 0.445 | 0.20 |
| **No QRS cleaning – Poor SNR subjects excluded** |  |  |  |  |  |
| **R-Ins Early** | 0.10 ± 5.33 | 2.01 ± 4.49 | -1.06 (19.3) | 0.302 | -0.39 |
| **R-Ins Late** | 2.56 ± 6.63 | 1.51 ± 5.33 | 0.47 (18.6) | 0.641 | 0.17 |
| **R-ACC Early** | 2.45 ± 7.68 | 0.80 ± 4.92 | 0.67 (15.9) | 0.510 | 0.26 |
| **R-ACC Late** | 1.73 ± 3.91 | -0.04 ± 5.15 | 1.14 (28.4) | 0.266 | 0.39 |
| **QRS cleaned - Poor SNR subjects excluded** |  |  |  |  |  |
| **R-Ins Early** | -0.65 ± 5.37 | -0.69 ± 4.87 | 0.02 (20.6) | 0.985 | 0.01 |
| **R-Ins Late** | 1.41 ± 7.65 | -1.32 ± 4.19 | 1.15 (14.5) | 0.269 | 0.44 |
| **R-ACC Early** | -5.62 ± 5.13 | -3.45 ± 3.50 | -1.32 (16.5) | 0.205 | -0.50 |
| **R-ACC Late** | 1.22 ± 3.79 | -0.74 ± 4.85 | 1.31 (27.7) | 0.200 | 0.45 |

##### 3.5.2.2. Hypertension (HTN vs NHTN)

###### Sensor space

No significant results were found in sensor space in either cohort, with or without QRS cleaning (table 6, figure 26, figure 27). When poor SNR subjects were excluded without QRS cleaning, a nominally significant early window emerged (p = 0.024, t(18.2) = -2.47, d = -1.02). This window, although attenuated, remained significant after QRS cleaning (p = 0.037, t(18.0) = -2.25, d = -0.93).

**Figure 25.**
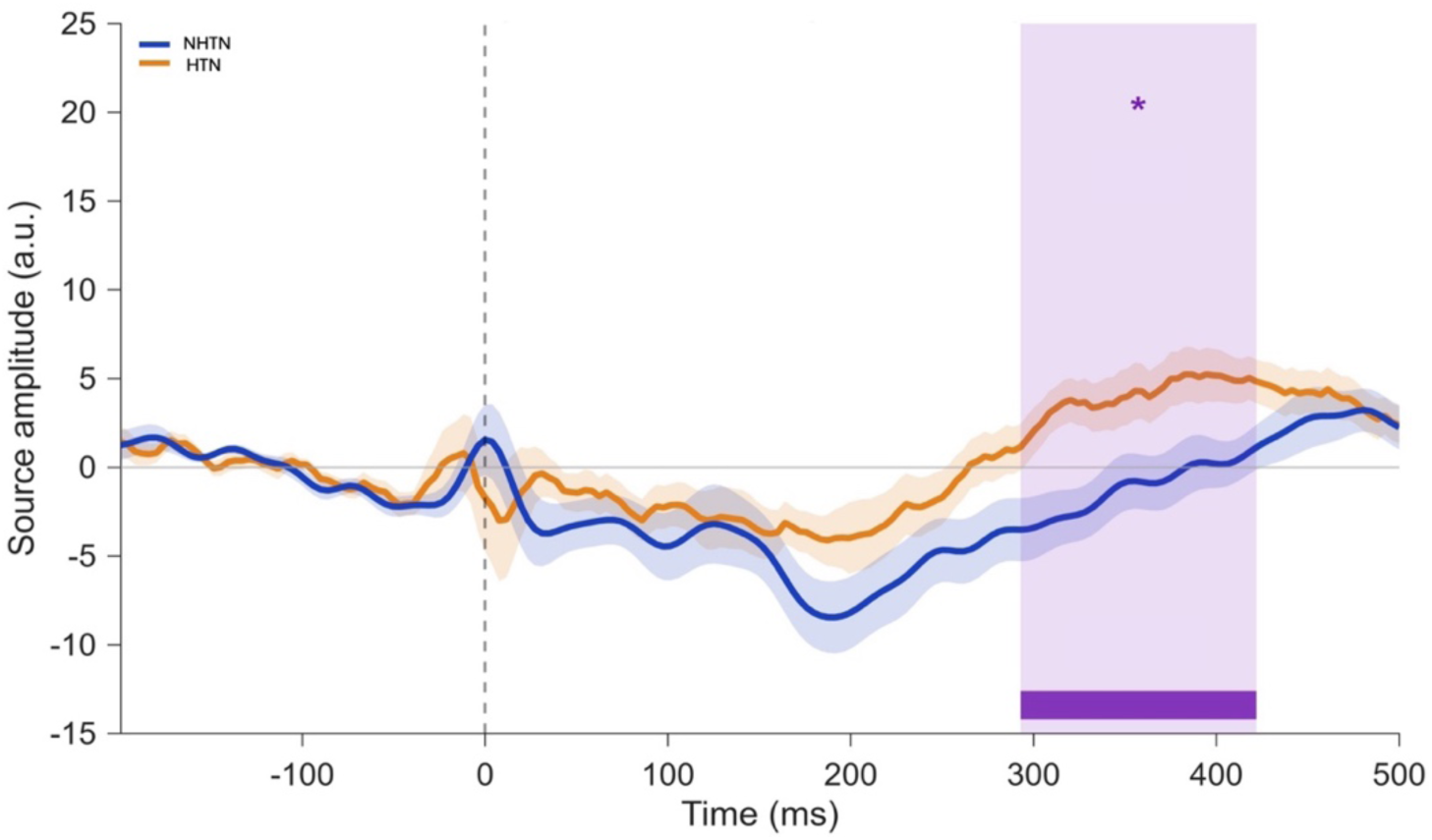
Group-averaged R-ACC source waveform for HTN vs NHTN subjects, without QRS cleaning. The purple shaded region denotes the significant cluster-corrected permutation window spanning 292-422 ms post R-peak (p = 0.019).

**Figure 26.**
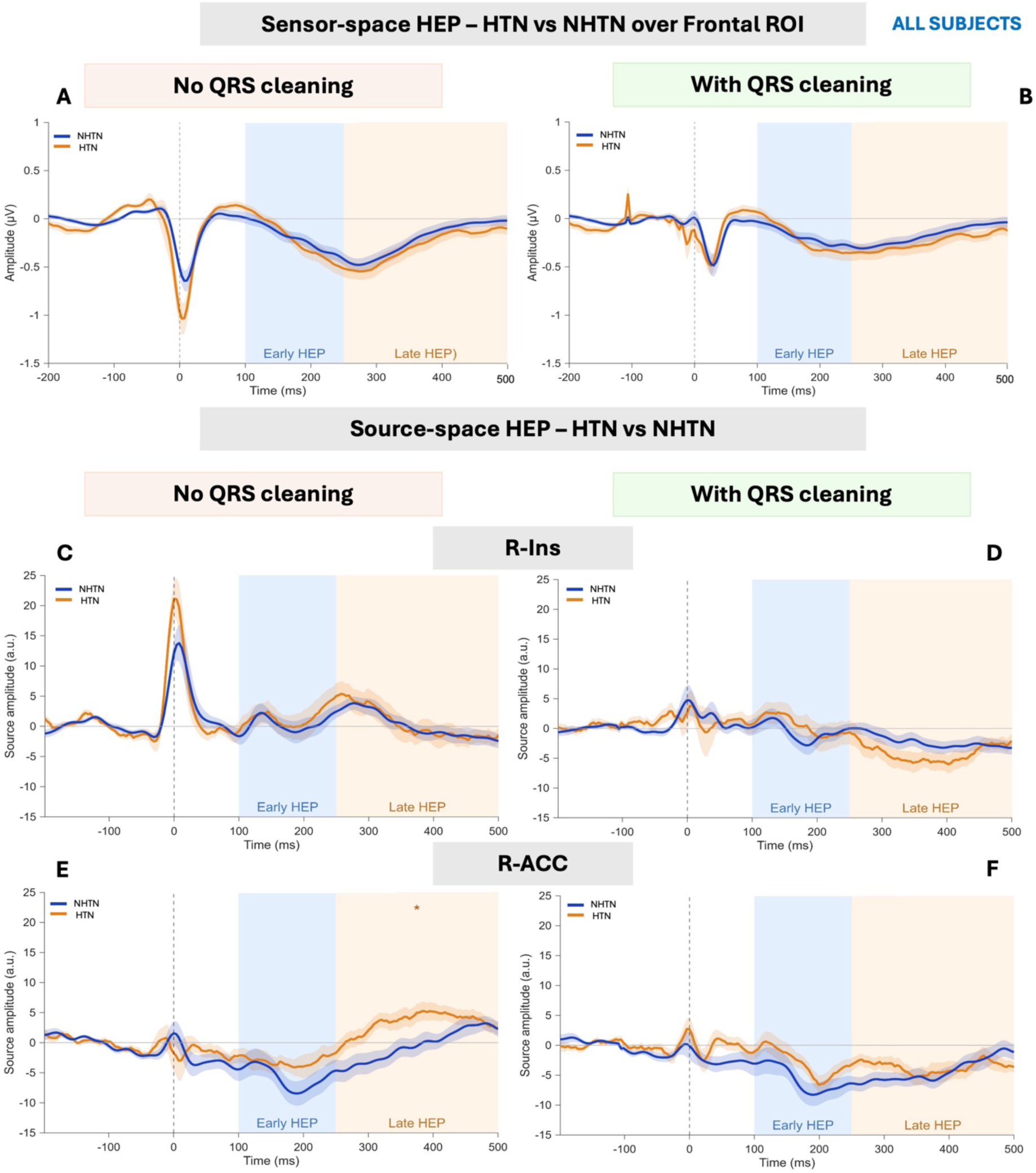
Grand average HEP in sensor space and source space before and after QRS cleaning across all subjects (HTN vs NHTN).

**Figure 27.**
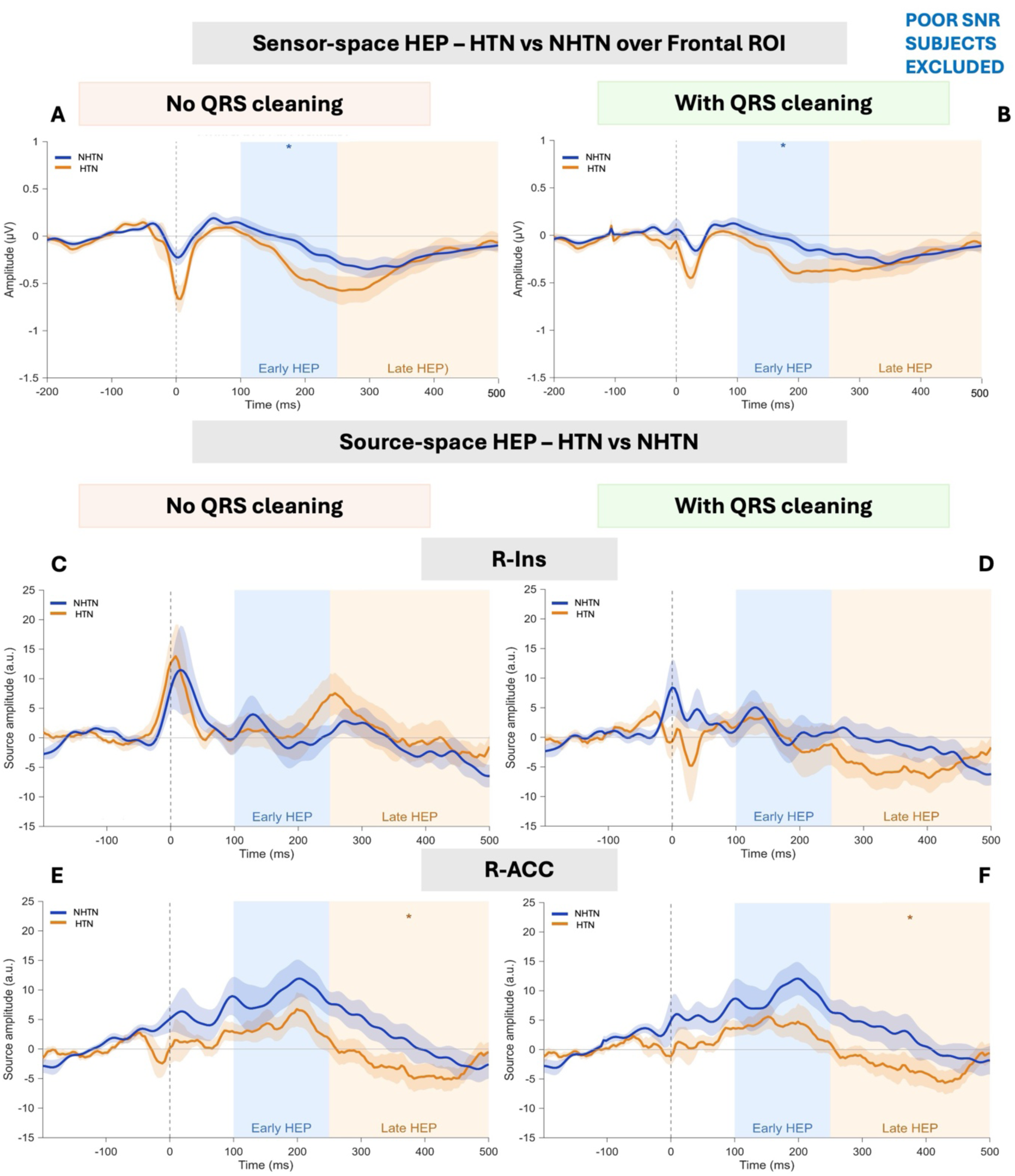
Grand average HEP in sensor space and source space before and after QRS cleaning when poor SNR subjects are excluded (HTN vs NHTN).

**Table 6.** Sensor-space HEP analysis HTN vs NHTN: mean frontal ROI amplitude and one-sample t-test statistics across pipelines and subject cohorts.

| Window | HTN<br>Mean ($\mu\text{V}$) + SD | NHTN<br>Mean ($\mu\text{V}$) + SD | t(df) | p | Cohen's d |
| --- | --- | --- | --- | --- | --- |
| **No QRS cleaning - All subjects** |  |  |  |  |  |
| **Early** | -0.21±0.27 | -0.20±0.27 | -0.17 (43.7) | 0.866 | -0.05 |
| **Late** | -0.28±0.33 | -0.21±0.26 | -0.81 (40.1) | 0.425 | -0.24 |
| **QRS cleaned - All subjects** |  |  |  |  |  |
| **Early** | -0.21±0.24 | -0.19±0.26 | -0.26 (44.0) | 0.796 | -0.08 |
| **Late** | -0.24±0.30 | -0.17±0.24 | -0.91 (40.4) | 0.370 | -0.27 |
| **No QRS cleaning – Poor SNR subjects excluded** |  |  |  |  |  |
| **Early** | **-0.29±0.28** | **-0.06±0.17** | **-2.47 (18.2)** | **0.024** | **-1.02** |
| **Late** | -0.30±0.40 | -0.23±0.23 | -0.46 (18.0) | 0.652 | -0.19 |
| **QRS cleaned - Poor SNR subjects excluded** |  |  |  |  |  |
| **Early** | **-0.25±0.27** | **-0.05±0.16** | **-2.25 (18.0)** | **0.037** | **-0.93** |
| **Late** | -0.25±0.35 | -0.21±0.22 | -0.36 (18.4) | 0.724 | -0.15 |

###### Source space

In source space without QRS cleaning (table 7), a significant HTN vs NHTN difference emerged in the R-ACC late window (p = 0.026; t(43.5) = 2.31, d = 0.68). Interestingly, this medium to large effect in source space, corroborated with cluster-corrected permutation testing, where a significant cluster spanning 292-422 ms post R-peak (p = 0.019) was identified (figure 25). This finding did not survive QRS cleaning (p=0.609; t(40.9) = 0.52, d = 0.15). All other VOIs and time windows were non-significant despite QRS cleaning.

**Table 7.** Source-space HEP analysis HTN vs NHTN: mean frontal ROI amplitude and one-sample t-test statistics across pipelines and subject cohorts.

| Window | HTN<br>Mean (a.u.) + SD | NHTN<br>Mean (a.u.) + SD | t(df) | p | Cohen's d |
| --- | --- | --- | --- | --- | --- |
| No QRS cleaning - All subjects |  |  |  |  |  |
| R-Ins Early | 1.47 ± 7.89 | 0.37 ± 5.73 | 0.54 (38.1) | 0.595 | 0.16 |
| R-Ins Late | 0.48 ± 7.45 | 0.24 ± 4.06 | 0.13 (31.8) | 0.895 | 0.04 |
| R-ACC Early | -3.08 ± 5.43 | -5.73 ± 8.51 | 1.27 (39.4) | 0.211 | 0.37 |
| R-ACC Late | 3.28 ± 5.05 | -0.53 ± 6.14 | 2.31 (43.5) | **0.026** | 0.68 |
| QRS cleaned - All subjects |  |  |  |  |  |
| R-Ins Early | -0.52 ± 7.54 | -0.52 ± 4.92 | 0.55 (35.6) | 0.587 | 0.16 |
| R-Ins Late | -4.08 ± 5.82 | -2.19 ± 3.43 | -1.33 (33.4) | 0.193 | -0.40 |
| R-ACC Early | -2.87 ± 5.18 | -5.65 ± 7.88 | 1.43 (40.1) | 0.161 | 0.42 |
| R-ACC Late | -3.62 ± 4.99 | -4.32 ± 4.12 | 0.52 (40.9) | 0.609 | 0.15 |
| No QRS cleaning - Poor SNR subjects excluded |  |  |  |  |  |
| R-Ins Early | 1.80 ± 10.19 | 0.50 ± 7.21 | 0.36 (19.8) | 0.726 | 0.15 |
| R-Ins Late | 5.78 ± 11.62 | 2.16 ± 6.48 | 0.93 (17.5) | 0.364 | 0.38 |
| R-ACC Early | 3.94 ± 6.07 | 9.17 ± 9.07 | -1.61 (17.3) | 0.126 | -0.68 |
| R-ACC Late | -0.06 ± 7.06 | 6.70 ± 7.67 | -2.19 (20.4) | **0.040** | 0.92 |
| QRS cleaned - Poor SNR subjects excluded |  |  |  |  |  |
| R-Ins Early | 0.42 ± 9.90 | 1.44 ± 6.08 | -0.30 (18.5) | 0.768 | -0.12 |
| R-Ins Late | -3.45 ± 9.58 | 0.82 ± 6.21 | -1.28 (19.0) | 0.217 | -0.53 |
| R-ACC Early | -3.78 ± 5.69 | -9.12 ± 8.20 | 1.80 (17.6) | 0.089 | 0.76 |
| R-ACC Late | 0.77 ± 5.30 | 5.58 ± 7.68 | 2.29 (17.6) | **0.035** | 0.96 |

With poor SNR subjects excluded in the absence of QRS cleaning, the late R-ACC significant effect resurfaced (p=0.040; t(20.4) = -2.19, d = 0.92). This effect survived and strengthened with QRS cleaning reflected by a large effect size (p = 0.035; t(17.6) = 2.29, d = 0.96). The R-Ins VOI showed no significant HTN vs NHTN differences across all pipelines.

#### 3.5.3. Cluster-corrected permutation test

For all source level comparisons and VOIs, a cluster-corrected permutation test was performed (5,000 permutations, cluster-forming threshold α=0.05). A significant cluster from 293 to 422 ms (p =0.019) (figure 25) was detected in the R-ACC VOI between all HTNs vs NHTNs subjects without QRS cleaning. This cluster falls within the late HEP window (250-500 ms post R-peak) and corroborates with R-ACC late window finding. No other comparison generated a significant cluster. The absence of significant clusters when poor SNR subjects are excluded could be due to reduced power resulting from the small group sizes (HTN n=12 and NHTN n=11).

### 3.6. Power analysis

Post-hoc power analysis was conducted for the source-space HTN vs NHTN R-ACC late HEP window comparison across each pipeline using a two-tailed independent-samples t-test (α = 0.05).

In the cohort including all subjects and in the absence of QRS cleaning (HTN n=22, NHTN n=24, d = 0.68), power reached 61.3%, demonstrating that the study was underpowered to reliably detect an effect of this magnitude. When poor SNR subjects were excluded, power was lower (55.4%), reflecting the trade-off between larder effect size (d = 0.92) and reduced sample sizes (HTN n=12, NHTN n=11). This was possibly due to the reduced sample sizes (HTN n=12, NHTN n=11). With QRS cleaning, achieved power was 59.5% (d = 0.96, HTN = 12, NHTN = 11). To achieve 80% power at the observed effect sizes, 36 subjects at minimum would be required for the full cohort. Approximately, 18-20 participants per group would be required when poor SNR subjects are excluded to reach 80% power. The large effect sizes (d = 0.68-0.96) indicate that a meaningful HEP is present, although the study is underpowered to reliably detect it on each run as excluding poor SNR subjects substantially reduces the sample size.

Power analyses revealed that the ANX vs NANX comparison was severely underpowered across the full cohort (30.2%, ANX n = 21, NANX n = 38), and when poor SNR subjects are excluded (18.4%, d<0.40). The consistently observed small effect sizes suggest that any true effect is likely negligible. However, larger sample sizes are required to explore this further.

## 4. Discussion

This study presents the first validated application of LCMV beamforming for HEP source extraction from EEG. Results demonstrate that this method can reliably recover HEP waveforms from deep interoceptive cortical regions in both simulated and clinical data. Across the three simulation models, beamforming achieved high waveform recovery fidelity under a wide range of SNR conditions, with near-perfect correlations (r>0.98 in model 1) achievable when source and sensor SNR were at moderate to high levels. The resilience of beamformer to CA is of central importance as CA has been the major obstacle in HEP research (Arnau et al., 2023; Dirlich et al., 1997) and appears to be continuously present throughout the epoch as evidence by figure 2. Although CA substantially increased inter-individual variability, mean waveform correlations remained robust at intermediate SNRs regardless of the model, confirming that spatial filtering in source space provides a means of separating cardiac and neural contributions that sensor-space (ICA-based) approaches cannot fully achieve (Buot et al., 2021; Petzschner et al., 2019).

A consistent finding across all simulated models was the asymmetric vulnerability of the R-Ins and R-ACC spatial filters to CA contamination. The insula filter was markedly affected by CA, as evidenced by greater waveform correlation degradation and increased localisation error compared to the ACC. This was particular to models 2 and 3 and could be explained by being a feature of the simulation. This finding has direct implications for the interpretation of HEP studies: insula-derived HEP estimates should be treated with caution when CA removal is incomplete, and cardiac signal null-projection should be applied as a post-processing step in analyses. In contrast, the ACC demonstrated superior spatial selectivity and greater resilience to CA across all models, suggesting it may be a more stable and reliable region for HEP source extraction.

Mapping empirical SNR estimates onto the simulation landscape demonstrated that stratification of empirical data against simulations provides a principled and data-driven approach for subject exclusion in HEP studies. Conventional exclusion criteria such as epoch rejection based on amplitude thresholds, minimum RR-intervals, operate at the data acquisition and preprocessing levels but are agnostic to downstream source-space estimates. The simulation-informed SNR mapping utilised here offers a complimentary metric. It allows to further filter empirical data by predicting source reconstruction reliability. Future work could expand this metric and increase the precision of SNR-mapping by expanding the simulation landscape with additional realistic noise models, and accounting for electrode placement, tissue conductivity and skull geometry amongst others.

Empirical results extended these insights with pooled results supporting the presence of a reliable HEP in both sensor and source space. After QRS cleaning, a significant R-Ins late window HEP effect emerged, previously absent without QRS cleaning. This follows the simulation results whereby CA seems to preferentially suppress R-Ins waveform recovery compared to R-ACC in a dual-source model. In contrast, the R-ACC demonstrated significant effects both with and without QRS cleaning across all subjects. QRS cleaning also eliminated a possible spurious late R-Ins window effect when poor SNR subjects were excluded. QRS cleaning would seem to suppress inflated results caused by residual CA and unmask true effects, suggesting that both beamforming and QRS cleaning are necessary steps in the pipeline to produce reliable and interpretable HEP waveforms.

Group differences detectable in source space appears to be absent in sensor-space. No significant results were found for HTN vs NHTN and for ANX vs NANX in sensor space across any pipeline, time window or patient cohort. Source space results revealed a significant late R-ACC difference between HTN and NHTN subjects in the full cohort without QRS cleaning (p = 0.026, d = 0.68), further supported by a significant cluster between 293-422 ms post R-peak (p = 0.019) extracted from cluster corrected permutation testing. When poor SNR subjects were excluded, this effect survived and strengthened with QRS cleaning. SNR-based data filtering and QRS cleaning to remove additional artefacts from CA would seem favourable steps to recover the most reliable source-level estimates. Significant effects not surviving QRS cleaning in the full cohort may reflect artefacts introduced by including poor SNR recordings, where CA contamination may also interact unpredictably with the beamforming output.

Additionally, none of the group differences observed in source space were detectable in sensor space analysis using a standardised HEP extraction pipeline. This highlights the added value of source-space analysis. The combination of spatial specificity (LCMV beamforming) and residual CA removal (QRS cleaning) appears to be both necessary and sufficient to reveal meaningful effects.

Because beamforming does not preserve the polarity of scalp-recorded signals, source waveforms are most meaningfully interpreted in terms of amplitude and localisation rather than the direction of their deflections. Interpreted on these terms, the LCMV beamforming results discussed here seem to align with recent work in the field (Gautier et al., 2025; Wang et al., 2025). Following Gautier et al. (2025) and Wang et al. (2025), an early (100-250 ms) and late (250-500 ms) HEP window post R-peak were examined, from which two prominent HEP components were recovered, corresponding to the early and delayed processing pathways proposed by Wang et al. (2025). In the empirical data, significant HEP activity was localised in the insula and ACC, in agreement with intracranial EEG evidence (Canales-Johnson et al., 2015; Park et al., 2018; Wang et al., 2025). Results across both regions exhibited heterogenous morphology and variable amplitude (Wang et al., 2025). The two-component temporal structure of HEP remains underexplored in the HEP literature. When investigated, the early and late responses have been inconsistently defined (Coll et al., 2021), with the early window sometimes being defined as 250-450 ms and the late window as 455-595 ms (Chi et al., 2026; Flasbeck et al., 2020), considerably later than the intervals adopted here. The functional significance of each components necessitates further investigation. Taken together, LCMV beamforming results suggest that there is a robust two-component temporal structure to HEP which warrants future work to resolve the timing of these windows and what they represent.

Beyond the findings discussed here, this work contributes a reusable and documented pipeline for applying LCMV beamforming to HEP extraction. Standardised source-level HEP pipelines are currently lacking, and existing HEP analyses in sensor space vary considerably in their processing choices. Preprocessing steps in sensor space prior to source reconstruction include exclusion of bad channels, filtering (0.5-30 Hz), epoching data around R-peaks, conversion to average reference, and baseline correction. A baseline correction window of -200 ms to -40 ms before the R-peak is recommended to avoid the QRS complex. By making this pipeline available within DAiSS (SPM release 2026 or later), we aim to make source-level analysis for HEP extraction more accessible and encourage methodological consistency across studies.

Several limitations of the present study warrant consideration. First, the simulation framework, while comprehensive, relied on fsaverage template head model. Empirical EEG data are acquired with individual head anatomies, electrode placements, different conductivity profiles that deviate from the template. Head model mismatch is a known source of localisation error in EEG beamforming (Dalal et al., 2014), and its effects are likely to be more pronounced for deep sources such as the insula than for superficial regions. Furthermore, spatial specificity remains uncertain without concurrent imaging, that is whether the reconstructed source waveforms originate specifically from the insula or ACC, rather than adjacent regions. Future work should evaluate the robustness of the pipeline to realistic forward model errors. Second, LCMV beamforming relied on the assumption that brain sources are temporally uncorrelated. This assumption could be partially violated in real resting-state EEG, where the insula and ACC activity may share common inputs from cardiac afferents and exhibit partial covariance. The cardiac signal null space projection approach, while shown to be effective, assumes that the CA subspace can be well characterised by subject specific ECG template and its temporal derivative. Some individuals may present with highly variable CA morphology, for example due to respiration, body movement or arrhythmias. The two-dimensional subspace may then be insufficient to fully capture the residual CA variance. In addition, to yield reliable beamforming results, a minimum of 64 channels is generally required. It is not recommended to interpolate channels as this could lead to rank-deficient covariance matrices (the rank is less than the number of channels) that would challenge the inversion step of the beamformer (Westner et al., 2022). Interpolating channels adds dimensions, i.e. channels but not new information, as interpolated channels are a linear combination of already existing channels. Finally, 55% of the empirical subjects met a high-SNR criterion with an expected correlation of r>0.40 based on the simulation landscape. This indicates that a substantial proportion of subjects yielded source waveforms of uncertain reliability. The SNR mapping offered an initial approximation to identify the SNR of empirical data in source and sensor space.

## 5. Conclusion

This study, the first of its kind, demonstrates that LCMV beamforming provides a valid and clinically informative approach to HEP source reconstruction from EEG. Simulation-based validation established the conditions under which reliable waveform recovery and accurate source localisation can be expected, providing a practical benchmark for future studies. Application of the LCMV beamforming pipeline to retrospective data revealed large and anatomically meaningful group differences in the right insula and right ACC in both anxiety and hypertension that were absent in conventional sensor-space analysis. The combination of LCMV beamforming with minimal prior-preprocessing, and post-hoc residual CA removal represents a coherent methodological framework for HEP source analysis that is generalisable across clinical populations and EEG system configurations. Future work should validate this pipeline with additional empirical data, explore its extension to the full interoceptive network, and investigate its sensitivity to within-subject variability in interoceptive processing.

## Data availability statement

the LCMV beamforming pipeline is publicly available on the SPM Development GitHub page within the DAiSS toolbox at spm/toolbox/DAiSS/bf_pipeline_LCMV_HEP_power_image.m and spm/toolbox/DAiSS/bf_pipeline_LCMV_HEP_source_extraction.m

## Funding statement

R.I.V is supported by the Biotechnology and Biological Sciences Research Council, BB/T008709/1. S.N.G is funded by a Wellcome Mental Health Award, 226778/Z/22/Z. M.Y. is funded MRC CARP award, MR/V037676/1.

## Conflict of interest disclosure

the authors attest to no competing interests.

## Ethics approval statement

this study did not involve testing on humans or animals.

## Notes

https://github.com/spm/spm/blob/main/toolbox/DAiSS/bf_pipeline_LCMV_HEP_power_image.m

https://github.com/spm/spm/blob/main/toolbox/DAiSS/bf_pipeline_LCMV_HEP_source_extraction.m

### Competing Interest Statement

The authors have declared no competing interest.

### Summary of Updates

This version of the manuscript has been revised to include the pipelines used for LCMV HEP beamforming now included in the DAiSS toolbox in SPM.

## References

Abolfathi, Y., Mohebbi, M., 2024. Heightened Heartbeat Evoked Potential in Obstructive Sleep Apnea Disorder During Sleep. IEEE Access 12, 189153–189162. 10.1109/ACCESS.2024.3439341

Arnau, S., Sharifian, F., Wascher, E., Larra, M.F., 2023. Removing the cardiac field artifact from the EEG using neural network regression. Psychophysiology 60, e14323. 10.1111/psyp.14323

Babo-Rebelo, M., Richter, C., Tallon-Baudry, C., 2016a. Neural Responses to Heartbeats in the Default Network Encode the Self in Spontaneous Thoughts. JOURNAL OF NEUROSCIENCE 36, 7829–7840. 10.1523/JNEUROSCI.0262-16.2016

Babo-Rebelo, M., Wolpert, N., Adam, C., Hasboun, D., Tallon-Baudry, C., 2016b. Is the cardiac monitoring function related to the self in both the default network and right anterior insula? PHILOSOPHICAL TRANSACTIONS OF THE ROYAL SOCIETY B-BIOLOGICAL SCIENCES 371. 10.1098/rstb.2016.0004

Baillet, S., Mosher, J.C., Leahy, R.M., 2001. Electromagnetic brain mapping. IEEE Signal Processing Magazine 18, 14–30. 10.1109/79.962275

Brookes, M.J., Mullinger, K.J., Stevenson, C.M., Morris, P.G., Bowtell, R., 2008. Simultaneous EEG source localisation and artifact rejection during concurrent fMRI by means of spatial filtering. Neuroimage 40, 1090–1104. 10.1016/j.neuroimage.2007.12.030

Buot, A., Azzalini, D., Chaumon, M., Tallon-Baudry, C., 2021. Does stroke volume influence heartbeat evoked responses? BIOLOGICAL PSYCHOLOGY 165. 10.1016/j.biopsycho.2021.108165

Canales-Johnson, A., Silva, C., Huepe, D., Rivera-Rei, A., Noreika, V., Garcia, M., Silva, W., Ciraolo, C., Vaucheret, E., Sedeno, L., Couto, B., Kargieman, L., Baglivo, F., Sigman, M., Chennu, S., Ibanez, A., Rodriguez, E., Bekinschtein, T., 2015. Auditory Feedback Differentially Modulates Behavioral and Neural Markers of Objective and Subjective Performance When Tapping to Your Heartbeat. CEREBRAL CORTEX 25, 4490–4503. 10.1093/cercor/bhv076

Chi, X., Hu, K., Ma, Y., Liu, Q., Chai, H., 2026. Impaired Processing of Bodily Signals Is Associated With Borderline Personality Traits: Insights From the Predictive Coding Model. Int J Psychol 61, e70178. 10.1002/ijop.70178

Coll, M., Hobson, H., Bird, G., Murphy, J., 2021. Systematic review and meta-analysis of the relationship between the heartbeat-evoked potential and interoception. NEUROSCIENCE AND BIOBEHAVIORAL REVIEWS 122, 190–200. 10.1016/j.neubiorev.2020.12.012

Dalal, S.S., Rampp, S., Willomitzer, F., Ettl, S., 2014. Consequences of EEG electrode position error on ultimate beamformer source reconstruction performance. Front. Neurosci. 8. 10.3389/fnins.2014.00042

Desmedt, O., Luminet, O., Walentynowicz, M., Corneille, O., 2023. The new measures of interoceptive accuracy: A systematic review and assessment. Neuroscience & Biobehavioral Reviews 153, 105388. 10.1016/j.neubiorev.2023.105388

Dirlich, G., Vogl, L., Plaschke, M., Strian, F., 1997. Cardiac field effects on the EEG. Electroencephalography and Clinical Neurophysiology 102, 307–315. 10.1016/S0013-4694(96)96506-2

Elkommos, S., Martin-Lopez, D., Koreki, A., Jolliffe, C., Kandasamy, R., Mula, M., Critchley, H.D., Edwards, M.J., Garfinkel, S., Richardson, M.P., Yogarajah, M., 2023. Changes in the heartbeat-evoked potential are associated with functional seizures. J Neurol Neurosurg Psychiatry 94, 769–775. 10.1136/jnnp-2022-330167

Flasbeck, V., Popkirov, S., Ebert, A., Brune, M., 2020. Altered interoception in patients with borderline personality disorder: a study using heartbeat-evoked potentials. BORDERLINE PERSONALITY DISORDER AND EMOTION DYSREGULATION 7. 10.1186/s40479-020-00139-1

Gautier, R., Latinus, M., Briend, F., 2025. Characterizing the Heartbeat-Evoked Potential: A Two-Component Model of Cardiac Signal Processing? Psychophysiology 62, e70206. 10.1111/psyp.70206

Henson, R.N., Abdulrahman, H., Flandin, G., Litvak, V., 2019. Multimodal Integration of M/EEG and f/MRI Data in SPM12. 10.3389/fnins.2019.00300

Hodossy, L., Ainley, V., Tsakiris, M., 2021. How do we relate to our heart? Neurobehavioral differences across three types of engagement with cardiac interoception. Biol Psychol 165, 108198. 10.1016/j.biopsycho.2021.108198

Kandasamy, R., Elkommos, S., van Rossum, I.A., Martin-Lopez, D., Koreki, A., Farrell, F., O’Sullivan, S., Diehl, B., Chowdhury, F.A., Critchley, H., Walker, M.C., Garfinkel, S., Yogarajah, M., 2026. The heartbeat evoked potential and the prediction of functional seizure semiology. Brain Commun 8, fcag120. 10.1093/braincomms/fcag120

Kern, M., Aertsen, A., Schulze-Bonhage, A., Ball, T., 2013. Heart cycle-related effects on event-related potentials, spectral power changes, and connectivity patterns in the human ECoG. NEUROIMAGE 81, 178–190. 10.1016/j.neuroimage.2013.05.042

Khalsa, S., Adolphs, R., Cameron, O., Critchley, H., Davenport, P., Feinstein, J., Feusner, J., Garfinkel, S., Lane, R., Mehling, W., Meuret, A., Nemeroff, C., Oppenheimer, S., Petzschner, F., Pollatos, O., Rhudy, J., Schramm, L., Simmons, W., Stein, M., Stephan, K., Van den Bergh, O., Van Diest, I., von Leupoldt, A., Paulus, M., Ainley, V., Al Zoubi, O., Aupperle, R., Avery, J., Baxter, L., Benke, C., Berner, L., Bodurka, J., Breese, E., Brown, T., Burrows, K., Cha, Y., Clausen, A., Cosgrove, K., Deville, D., Duncan, L., Duquette, P., Ekhtiari, H., Fine, T., Ford, B., Cordero, I., Gleghorn, D., Guereca, Y., Harrison, N., Hassanpour, M., Hechler, T., Heller, A., Hellman, N., Herbert, B., Jarrahi, B., Kerr, K., Kirlic, N., Klabunde, M., Kraynak, T., Kriegsman, M., Kroll, J., Kuplicki, R., Lapidus, R., Le, T., Hagen, K., Mayeli, A., Morris, A., Naqvi, N., Oldroyd, K., Pane-Farre, C., Phillips, R., Poppa, T., Potter, W., Puhl, M., Safron, A., Sala, M., Savitz, J., Saxon, H., Schoenhals, W., Stanwell-Smith, C., Teed, A., Terasawa, Y., Thompson, K., Toups, M., Umeda, S., Upshaw, V., Victor, T., Wierenga, C., Wohlrab, C., Yeh, H., Yoris, A., Zeidan, F., Zotev, V., Zucker, N., Interoception Summit 2016 Particip, 2018. Interoception and Mental Health: A Roadmap. BIOLOGICAL PSYCHIATRY-COGNITIVE NEUROSCIENCE AND NEUROIMAGING 3, 501–513. 10.1016/j.bpsc.2017.12.004

Litvak, V., Mattout, J., Kiebel, S., Phillips, C., Henson, R., Kilner, J., Barnes, G., Oostenveld, R., Daunizeau, J., Flandin, G., Penny, W., Friston, K., 2011. EEG and MEG Data Analysis in SPM8. Comput Intell Neurosci 2011, 852961. 10.1155/2011/852961

Obeid C Picone, 2016. The Temple University Hospital EEG data corpus. 10.34944/dspace/5136

Pang, J., Tang, X., Li, H., Hu, Q., Cui, H., Zhang, L., Li, W., Zhu, Z., Wang, J., Li, C., 2019. Altered Interoceptive Processing in Generalized Anxiety Disorder A Heartbeat - Evoked Potential Research. FRONTIERS IN PSYCHIATRY 10. 10.3389/fpsyt.2019.00616

Park, H., Blanke, O., 2019. Heartbeat-evoked cortical responses: Underlying mechanisms, functional roles, and methodological considerations. NEUROIMAGE 197, 502–511. 10.1016/j.neuroimage.2019.04.081

Park, H.-D., Bernasconi, F., Salomon, R., Tallon-Baudry, C., Spinelli, L., Seeck, M., Schaller, K., Blanke, O., 2018. Neural Sources and Underlying Mechanisms of Neural Responses to Heartbeats, and their Role in Bodily Self-consciousness: An Intracranial EEG Study. Cereb Cortex 28, 2351–2364. 10.1093/cercor/bhx136

Park, H.-D., Blanke, O., 2019. Heartbeat-evoked cortical responses: Underlying mechanisms, functional roles, and methodological considerations. Neuroimage 197, 502–511. 10.1016/j.neuroimage.2019.04.081

Park, H.-D., Correia, S., Ducorps, A., Tallon-Baudry, C., 2014. Spontaneous fluctuations in neural responses to heartbeats predict visual detection. Nat Neurosci 17, 612–618. 10.1038/nn.3671

Petzschner, F.H., Weber, L.A., Wellstein, K.V., Paolini, G., Do, C.T., Stephan, K.E., 2019. Focus of attention modulates the heartbeat evoked potential. Neuroimage 186, 595–606. 10.1016/j.neuroimage.2018.11.037

Petzsehner, F., Weber, L., Wellstein, K., Paolini, G., Do, C., Stephan, K., 2019. Focus of attention modulates the heartbeat evoked potential. NEUROIMAGE 186, 595– 606. 10.1016/j.neuroimage.2018.11.037

Pollatos, O., Schandry, R., 2004. Accuracy of heartbeat perception is reflected in the amplitude of the heartbeat-evoked brain potential. PSYCHOPHYSIOLOGY 41, 476–482. 10.1111/1469-8986.2004.00170.x

Schandry, R., Sparrer, B., Weitkunat, R., 1986. From the heart to the brain: A study of heartbeat contingent scalp potentials. International Journal of Neuroscience 30, 261–275. 10.3109/00207458608985677

Suksasilp, C., Garfinkel, S.N., 2022. Towards a comprehensive assessment of interoception in a multi-dimensional framework. Biol Psychol 168, 108262. 10.1016/j.biopsycho.2022.108262

Van Veen, B.D., Buckley, K.M., 1988. Beamforming: a versatile approach to spatial filtering. IEEE ASSP Mag. 5, 4–24. 10.1109/53.665

Van Veen, B.D., Van Drongelen, W., Yuchtman, M., Suzuki, A., 1997. Localization of brain electrical activity via linearly constrained minimum variance spatial filtering. IEEE Transactions on Biomedical Engineering 44, 867–880. 10.1109/10.623056

Virjee, R.-I., Kandasamy, R., Garfinkel, S.N., Carmichael, D.W., Yogarajah, M., 2026. Review of methods to derive the heartbeat-evoked potential: past practices and future directions. Soc Cogn Affect Neurosci nsag057. 10.1093/scan/nsag057

Wang, X., Yang, H., Cheng, Y., Liu, S., Jin, G., Qiao, Z., Qi, L., Wang, S., Ge, J., Hu, D., Tang, H., Gao, R., Xu, C., Zhang, X., Wang, D., Xue, X., Dai, A., Zhao, W., Yu, T., Wang, Y., Si, B., Zhao, G., Ren, L., 2025. Mapping human brain topography to heart rhythms: an SEEG study. Cardiovasc Res 121, 1228–1239. 10.1093/cvr/cvaf099

Westner, B.U., Dalal, S.S., Gramfort, A., Litvak, V., Mosher, J.C., Oostenveld, R., Schoffelen, J.-M., 2022. A unified view on beamformers for M/EEG source reconstruction. NeuroImage 246, 118789. 10.1016/j.neuroimage.2021.118789

Woolrich, M., Hunt, L., Groves, A., Barnes, G., 2011. MEG Beamforming using Bayesian PCA for Adaptive Data Covariance Matrix Regularisation. Neuroimage 57, 1466–1479. 10.1016/j.neuroimage.2011.04.041

Yoris, A., Abrevaya, S., Esteves, S., Salamone, P., Lori, N., Martorell, M., Legaz, A., Alifano, F., Petroni, A., Sanchez, R., Sedeno, L., Garcia, A., Ibanez, A., 2018. Multilevel convergence of interoceptive impairments in hypertension: New evidence of disrupted body-brain interactions. HUMAN BRAIN MAPPING 39, 1563–1581. 10.1002/hbm.23933

Yoris, A., García, A.M., Traiber, L., Santamaría-García, H., Martorell, M., Alifano, F., Kichic, R., Moser, J.S., Cetkovich, M., Manes, F., Ibáñez, A., Sedeño, L., 2017. The inner world of overactive monitoring: neural markers of interoception in obsessive–compulsive disorder. Psychological Medicine 47, 1957–1970. 10.1017/S0033291717000368

